# Blockade of TGF-β signaling reactivates HIV-1/SIV reservoirs and immune responses *in vivo*

**DOI:** 10.1101/2022.05.13.489595

**Authors:** S Samer, Y Thomas, M Araínga, CM Carter, LM Shirreff, MS Arif, JM Avita, I Frank, M McRaven, CT Thuruthiyil, V Heybeli, MR Anderson, B Owen, A Gaisin, D Bose, LM Simons, JF Hultquist, J Arthos, C Cicala, I Sereti, P Santangelo, R Lorenzo-Redondo, TJ Hope, FJ Villinger, E Martinelli

## Abstract

Elevated levels of TGF-β, a potent immunosuppressive factor, are present in HIV-1 infected individuals even after years of antiretroviral therapy (ART). TGF-β plays a critical role in maintaining immune cells in a resting state by inhibiting cell activation and proliferation. Resting HIV-1 target cells represent one of the main cellular reservoirs after long term ART and the low inducibility of the latent provirus constitutes one of the major obstacles to “kick and kill” cure strategies. We hypothesized that releasing cells from TGF-β-driven signaling would promote latency reversal. To test our hypothesis, we compared *ex vivo* models of HIV-1 latency reactivation with and without TGF-β and a TGF-β type 1 receptor (TGFBR1) inhibitor, galunisertib. We also tested the effect of galunisertib in SIV infected, ART treated macaques by monitoring SIV envelope (env) protein expression via PET/CT using the Cu^64^-anti gp120 Fab (7D3) probe, along with plasma and tissue viral loads (VL). Exogenous TGF-1β reduced HIV-1 reactivation in U1 and ACH2 latency models. Galunisertib increased HIV-1 latency reversal both in *ex vivo* models and in PBMC from HIV-1 infected, cART treated aviremic donors. *In vivo*, oral galunisertib promoted increased SIV env protein total standardized uptake values (SUVtot) in PET/CT images of tissues (gut and lymph nodes) of 5 out of 7 aviremic, long-term ART-treated, SIV-infected, macaques. This increase correlated with an increase in SIV RNA in gut tissue. Two out of 7 animals also exhibited increases in plasma viral load. Higher anti-SIV T cell responses and anti-SIV env antibody titers were detected after galunisertib treatment in most animals. In summary, our data suggest that blocking TGF-β signaling simultaneously increases retroviral reactivation events and enhances anti-SIV immune responses.

## Introduction

Despite the success of combination ART (ART) in suppressing viral replication and preventing immunodeficiency progression, HIV-1 infection persists and evades immune responses in viral reservoirs, which are considered the major obstacle to HIV-1 cure (*1-3*). Long-lived CD4^+^ T cells that expand by homeostatic or antigen-driven proliferation in blood and tissues constitute a reservoir of proviruses that may be fully latent or transcriptionally active to a different extent in different CD4^+^ T cells(*4-7*). Myeloid cells may also constitute important cellular reservoir(*8*), although HIV-1 latency in the myeloid compartment has only recently been explored(*8-10*). Viral rebound upon ART interruption occurs in the vast majority of patients, likely following stochastic HIV-1 reactivation events and its dynamics depend on the size of the reservoir and host immune factors(*11*) (*12-15*). “Shock and kill” strategies have the potential to lead to a functional HIV-1 cure by reducing reservoir size and stimulating immune responses(*16, 17*). However, the success of this strategy is hindered by several obstacles including the vast heterogeneity of the cells comprising the reservoir and of the mechanisms of latency(*4, 17-19*) and the low inducibility of the latent proviruses(*20, 21*).

TGF-β is one of the most potent endogenous immunosuppressive factors and it is considered the master regulator of mucosal immunity(*22-24*). TGF-β signaling regulates diverse cellular processes, including proliferation, differentiation, and migration(*25*), and influences developmental as well as pathological processes such as epithelial to mesenchymal (EMT) transition in fibrosis(*26*). In the immune system, TGF-β is essential to the maintenance of tolerance(*27-29*). Importantly, TGF-β is the most prominent factor responsible for the maintenance of a resting state in CD4^+^ T cells(*30*) and loss of TGF-β responsiveness in mature T cells decreases their threshold for activation(*31*). Moreover, TGF-β also impacts activation status and function of myeloid cells by decreasing antigen presentation capability and maturation and inducing tolerogenic properties(*29, 32, 33*).

Since the beginning of the HIV-1 epidemic several reports have documented higher concentrations of TGF-β in the blood, lymphoid tissues, and cerebrospinal fluid of people living with HIV-1 (PLWH)(*34-36*). Higher levels of TGF-β in PLWH and SIV-infected macaques have been linked to disease progression(*34*) (*37, 38*). PBMCs from PLWH spontaneously release high levels of TGF-β (*39, 40*) (*41, 42*). Among other mechanisms, TGF-β release particularly in the gut tissue is the result of a biological anti-inflammatory process counteracting HIV-driven microbial translocation and chronic inflammation(*43*). Paradoxically, this anti-inflammatory, TGF-β-centered response not only exacerbates immunosuppression, but also predisposes for the development of non-AIDS-related, non-communicable disorders(*43*). Finally, TGF-β-driven fibrosis of lymphoid tissues has been implicated in the CD4^+^ T cell loss before ART initiation(*44, 45*) and failure to achieve full immune reconstitution with ART(*46*). Importantly, TGF-β inhibits TCR-induced T cell proliferation, CD28-mediated co-stimulation, and activation of T cells (*47-49*). Indeed, it has recently been used to promote latency in *ex vivo* models with primary polarized effector CD4^+^ T cells (*50-52*). Moreover, TGF-β was shown to induce latency in other cell types susceptible to HIV infection(*53*).

The 3 isoforms of TGF-β (TGF-β1, TGF-β2 and TGF-β3) bind to the specific type I receptor TGFBR1/ALK5 which then undergoes dimerization with the type 2 TGF-βRII. The differential function and signaling mechanism of these major isoforms is still under investigation. However, TGF-β1 appears to be the most prominent regulator of immune responses(*54-56*). The TGF-β receptor complex then signals through a canonical pathway by phosphorylating Smad2 and/or Smad3. The Smad complex formed by co-heterodimers with Smad4 interacts with a large variety of co-activators and co-repressor in the nucleus and regulates the transcription of hundreds of genes in a cell type-specific, context dependent way(*29*). Galunisertib (LY2157299), a small-molecule selective inhibitor of the TGFBR1/ALK5 (*57, 58*), has been developed by Eli Lilly as an anti-cancer therapeutic to target the direct and indirect effect of TGF-β on tumor growth. Galunisertib (Gal) progressed to an advanced stage of clinical development (Phase II) and had been tested in several different trials for different solid tumors(*58-61*), when Eli Lilly decided to end its program to pursue more promising targets(*62*).

Given the suppressive activity of TGF-β on different immune cell targets of HIV-1 infection and its role in latency induction(*50, 53*), we hypothesized that TGF-β may also exert a suppressive effect on latency reactivation. Moreover, we hypothesized that if TGF-β inhibited TCR- or PKC-driven HIV reactivation from latency, inhibition of TGF-β signaling may, in turn, increase the frequency of latency reversal following stimulation of these pathways. We tested this hypothesis using *ex vivo* models of HIV latency including PBMC from aviremic, ART treated PLWH. Finally, we tested the impact of the TGFBR1 inhibitor galunisertib *ex vivo* and in SIV infected ART treated macaques. Our data suggest that targeting TGF-β signaling may constitute a novel way to increase viral latency reactivation during ART while potentiating antiviral immune responses.

## Results

### TGF-β1 inhibits PMA-driven and DC-driven HIV-1 latency reactivation *in vitro*

We tested the impact of exogenous TGF-β1 on phorbol 12-myristate 13 acetate-(PMA)-induced HIV reactivation in the U1 and ACH-2 classical cell line models of HIV latency (U1 derived from U937 promonocytic cells(*63*) and ACH-2 derived from CEM T cells(*64*)). U1 cells originally contained 2 copies of integrated HIV DNA, although more recently HIV was detected in additional sites(*65*) and latency is maintained by mutations in the Tat protein(*63*) (*66*). The frequency of U1 cells expressing p24-Gag at baseline is very low (Fig 1A, MOCK-DMSO), but it is readily induced by stimulation with PMA (Fig 1A MOCK-PMA). Addition of 10ng/ml of TGF-β1 to the culturing media decreases the frequency of p24-gag+ cells below the baseline level in unstimulated cultures and reduces the PMA-driven increase in the frequency of p24-gag+ cells by ∼20% (Fig 1A, B and D). In contrast, blockade of TGF-β1 signaling using galunisertib restores the frequency of p24-gag+ cells in the presence of exogenous TGF-β1 to baseline or PMA-induced levels. Treatment with 1μM of galunisertib alone appears to slightly increase the frequency of p24-gag+ cells from baseline but has no impact on PMA-stimulated U1 cells (Fig 1D).

**Figure 1.**
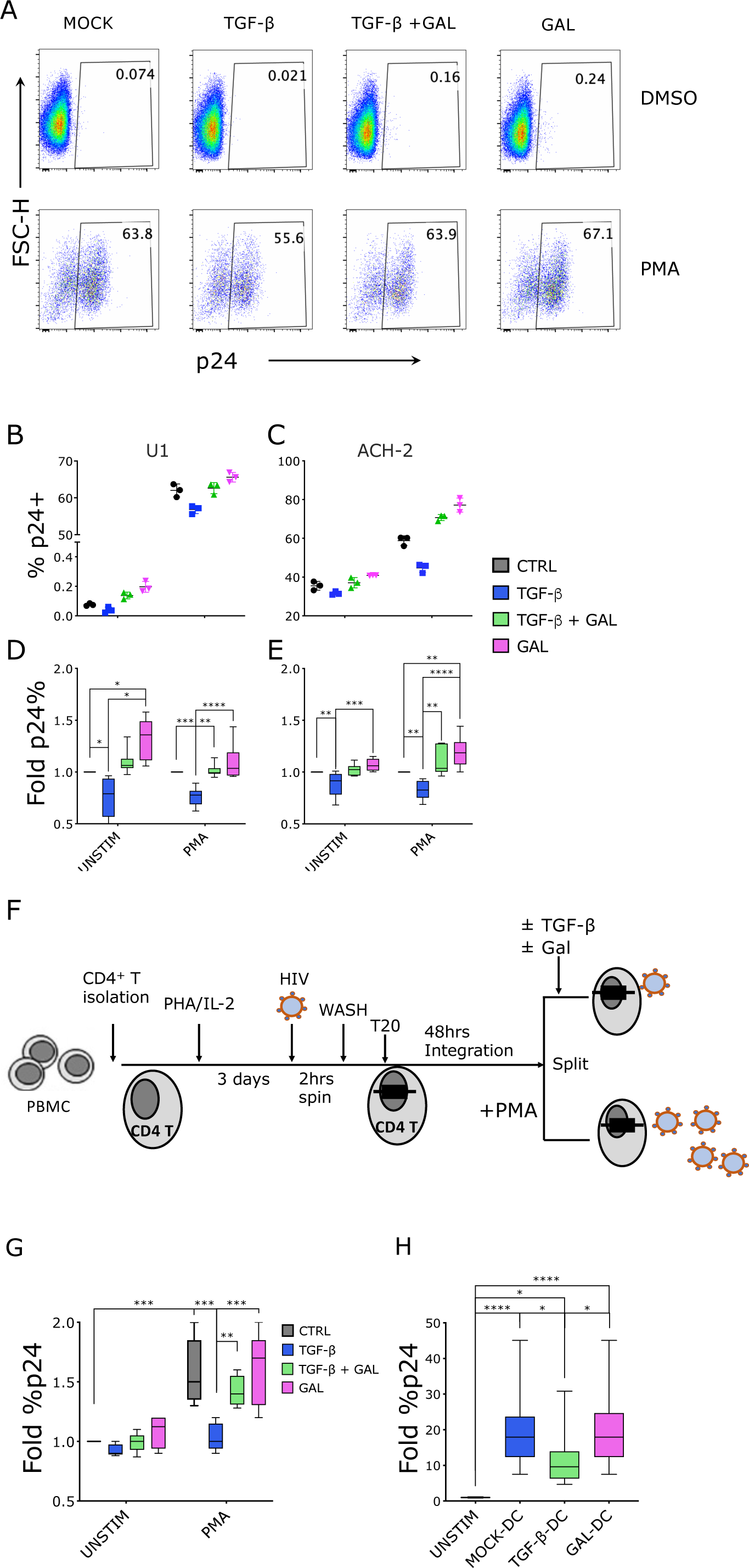
TGF-β inhibits HIV-1 latency reactivation *in vitro*. **A-E)** U1 and ACH-2 cells were treated with TGF-β1 (10ng/ml) or galunisertib (1uM), both or mock treated in presence vs absence of PMA (100ng/ml) for 18hrs and then stained for intracellular p24. A) An example of flow cytometry data for p24 detection in one of the experiments with U1 cells. B-C) Raw p24 data from one representative experiment with U1 (B) or ACH-2 (C) cells (black circles: mock; blue squares: TGF-β1; green triangles: TGF-β1+ gal; pink triangles: Gal-only). D-E) Summary data of fold increase in the frequency of p24+ cells over the mock condition (box plot with median and min max whiskers) are shown from 5 similar experiments (U1 on the left; ACH-2 on the right). F-G) For this primary CD4^+^ T cell model of latency, CD4^+^ T cells were isolated from PBMC, activated, infected by spinoculation and incubated for 2 days in presence of T20. PMA (100ng/ml) was used for reactivation of latently infected cells for 18hrs in presence of TGF-β1 (10ng/ml) or galunisertib (1uM), both or mock treatment. A schematic of the experiment is shown in F). G) A summary of the data (fold increase over the unstimulated condition) from 5 experiments with cells from different donors run in triplicate (box plot with median and min max whiskers). H) Data from a model of DC-driven HIV reactivation from U1 cells are shown. Fold difference in the frequency of p24+ U1 cells in absence vs presence of moDCs or TGF-β DCs are shown for 5 different experiments in triplicate (box plot with median and min max whiskers). A-H) Conditions were compared by Kruskal-Wallis ANOVA test followed by the Dunn’s test corrected for multiple comparisons (Significant *p*-values of α<0.05 (*), α<0.01 (**) and α <0.001 (***) are indicated).

ACH-2 cells were originally reported to carry a single copy of HIV DNA(*64*), but, as with U1, HIV integration has been detected in multiple sites in more recent culture passages(*65*). Latency is maintained by a mutation in TAR that impairs the LTR response to Tat(*67*). In contrast to U1, the frequency of p24-gag+ cells is relatively high in ACH-2 at baseline (Fig S1A, MOCK-DMSO), but increases further in response to PMA (Fig S1A, MOCK-PMA). We found that TGF-β1 was able to decrease the frequency of p24-gag+ cells in PMA-stimulated cultures by ∼20% and a smaller but consistent decrease in baseline p24-gag levels was also detected in absence of PMA (Fig 1C and E). Galunisertib treatment was able to restore frequency of p24-gag+ cells in TGF-β1-treated cultures and increase it above PMA-stimulated cultures (Fig 1C and E) when administered in absence of exogenous TGF-β1. This increase may be attributed to blockade of TGF-β1 in the fetal bovine serum (FBS) used to supplement the culturing media(*30*).

Next, we tested the impact of TGF-β1 on a primary model of HIV-1 latency using freshly isolated CD4^+^ T cells. The experiment was based on the model by Berkhout and colleagues(*68*) with some modifications (Fig 1F). Specifically, we isolated CD4^+^ T cells from HIV-1 uninfected, de-identified donor PBMC and activated them with PHA and IL-2 for 3 days before infection via spinoculation with HIV ADA. After extensive wash of the viral inoculum, cells were cultured 48hrs in presence of T20 (enfuvirtide) to avoid viral spread beyond the cells with initial integrations. The overall initial infection achieved was very low despite the spinoculation step (∼0.1% of p24-gag+ cells, Fig S2). However, PMA stimulation was able to increase the frequency of p24-gag+ cells consistently (Fig 1G) and this was significantly inhibited when TGF-β1 (10ng/ml) was added concomitantly with PMA. In contrast the addition of both TGF-β1 and galunisertib (1μM) or galunisertib alone did not alter the frequency of p24-gag+ cells from the level in control cultures (Fig 1G).

Finally, considering that dendritic cells (DC) are important players in reversing HIV latency(*69*) and that TGF-β1 impairs DC activation(*70, 71*), we decided to test the impact of TGF-β1 on DC-mediated HIV-1 latency reactivation(*68, 72*). Monocyte derived (mo)DCs were generated from primary monocytes as previously described(*73*) and treated or not with TGF-β1 for the last 2 days of culture. moDCs generated in the presence of TGF-β1 exhibited a less mature phenotype with tendency toward lower HLA-DR and significantly lower DEC-205 expression (Fig S3A). When co-cultured with eFluo670 pre-labeled U1 cells, moDCs were able to promptly activate HIV-1 expression with the frequency of p24-Gag+ cells increasing of several folds within a few hours (Fig 1H and S3B). However, the frequency of p24-gag+ cells in the cocultures with TGF-β1-treated DCs was significantly less than in co-cultures with untreated DCs and this was blocked when TGF-β1-treatment of DCs occurred in presence of galunisertib (Fig 1H and S3B).

### Blocking TGF-β1 signaling increases LRA-induced HIV-1 reactivation *ex vivo*

Considering that TGF-β is present in FBS, used to supplement PBMC and T cell cultures, and that PBMC from people living with HIV-1 release TGF-β in culture supernatants(*39, 74, 75*), we hypothesized that blocking TGF-β while reversing HIV-1 latency *ex vivo* from PBMC of ART-treated, aviremic individuals may lead to higher frequency of HIV-1 reactivation than in untreated cultures. To test this hypothesis, we used a modified version of the classical quantitative viral outgrowth assay (qVOA)(*76*). We used whole PBMC to assess potential latency reactivation from any HIV-1-infected circulating cell and not only from resting CD4^+^ T cells (Fig 2A). PBMC from two different sets of aviremic, ART-treated donors (Table 1) were activated with PMA (at 100ng/ml or 10ng/ml) and with Vorinostat (SAHA, at 1μM) in presence of T20 and galunisertib (1μM) or mock solution. After overnight incubation supernatants were collected and assayed for both vRNA in a subset of donors (n=5) and viral outgrowth (n=9). After washing out the PMA, cells were plated at 3×10^5^/well in 96 well plates with or without galunisertib in replicates (>14 replicate wells) and SupT1 cells were added to amplify the infection. Infectious unit per million cells (IUPM) were calculated based on the number of infected wells as determined by p24 ELISA (Fig 2A). A significantly higher frequency of latency reactivation events occurred in presence of galunisertib in PBMC from most donors (Fig 2B). This occurred with both high and low PMA concentrations and when reactivation was performed with Vorinostat. No other latency reversal agents (LRA) were tested. Moreover, higher levels of vRNA were detected after PMA stimulation in galunisertib - treated PBMC than in mock treated (Fig 2C).

**Table 1.**
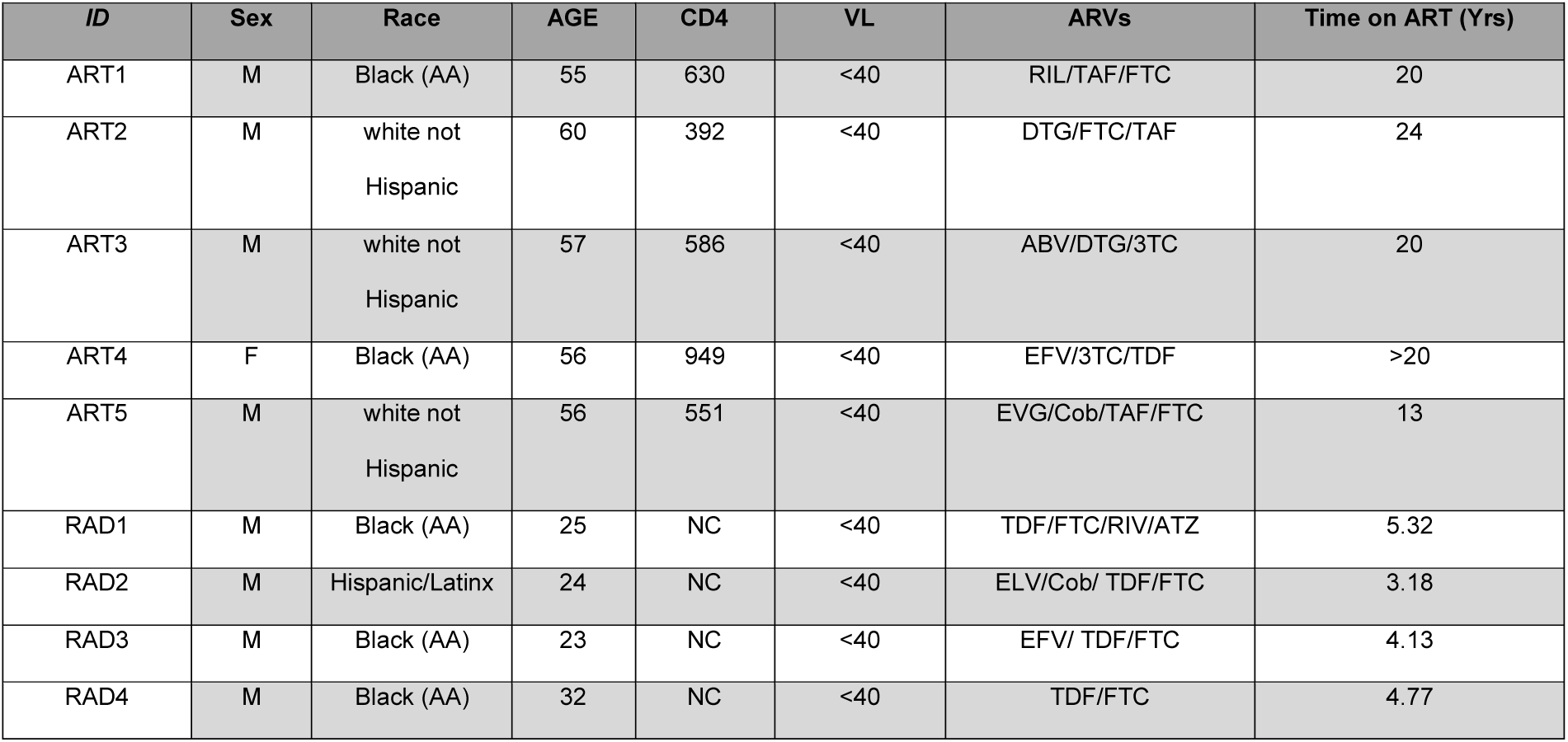
Participants characteristics.

| *ID* | *Sex* | *Race* | *AGE* | *CD4* | *VL* | *ARVs* | *Time on ART (Yrs)* |
| --- | --- | --- | --- | --- | --- | --- | --- |
| ART1 | M | Black (AA) | 55 | 630 | <40 | RIL/TAF/FTC | 20 |
| ART2 | M | white not<br>Hispanic | 60 | 392 | <40 | DTG/FTC/TAF | 24 |
| ART3 | M | white not<br>Hispanic | 57 | 586 | <40 | ABV/DTG/3TC | 20 |
| ART4 | F | Black (AA) | 56 | 949 | <40 | EFV/3TC/TDF | >20 |
| ART5 | M | white not<br>Hispanic | 56 | 551 | <40 | EVG/Cob/TAF/FTC | 13 |
| RAD1 | M | Black (AA) | 25 | NC | <40 | TDF/FTC/RIV/ATZ | 5.32 |
| RAD2 | M | Hispanic/Latinx | 24 | NC | <40 | ELV/Cob/ TDF/FTC | 3.18 |
| RAD3 | M | Black (AA) | 23 | NC | <40 | EFV/ TDF/FTC | 4.13 |
| RAD4 | M | Black (AA) | 32 | NC | <40 | TDF/FTC | 4.77 |

**Figure 2.**
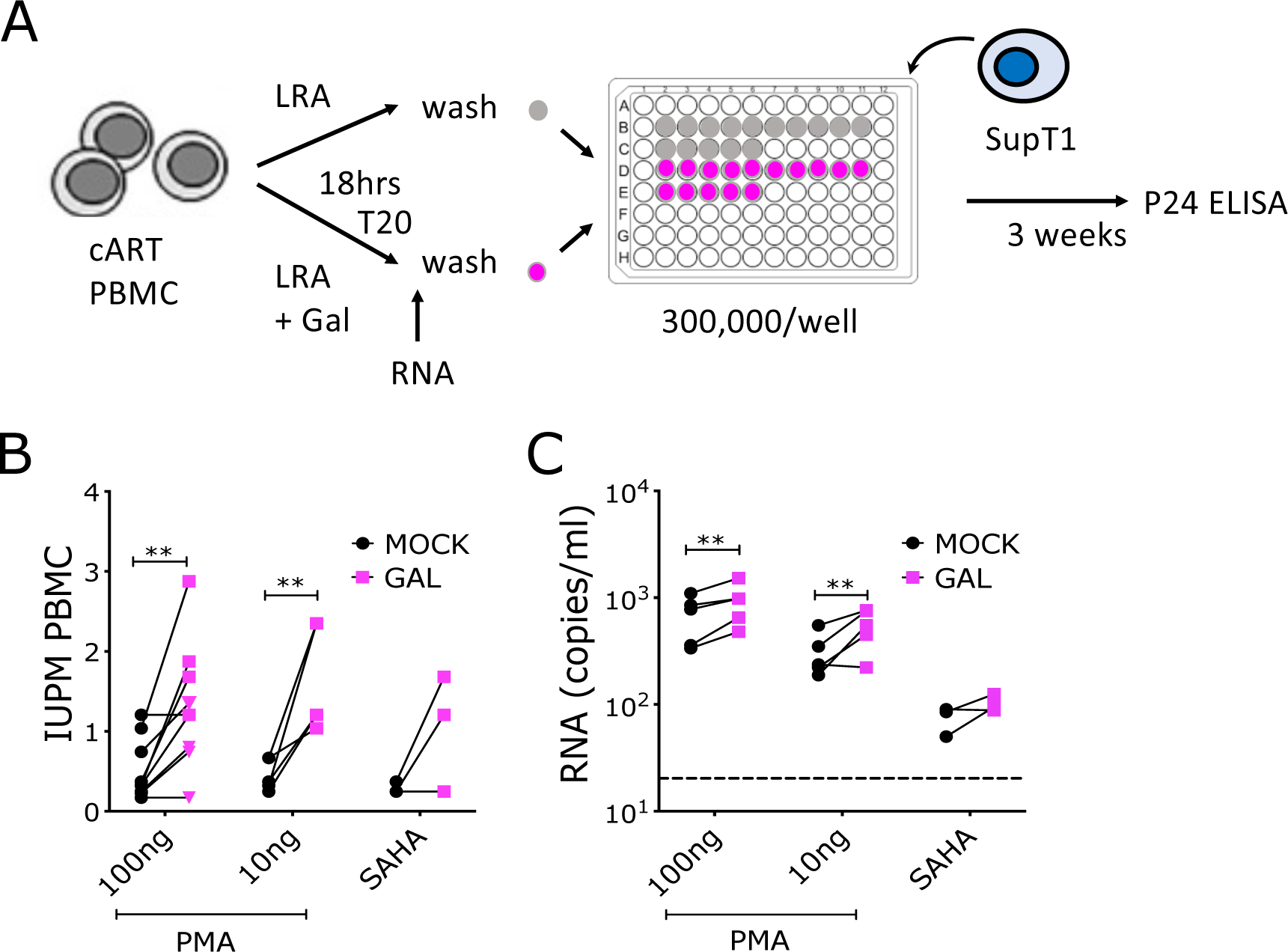
Blocking TGF-β1 signaling increases LRA-induced HIV-1 reactivation *ex vivo*. De-identified PBMC from aviremic, ARV-treated PLWH collected at NIH and NU were used for viral outgrowth assays to estimate the frequency of HIV infected cells able to produce replication competent virus within the unfractionated PBMC. A) Schematic of the qVOA assay where activation with a latency reversal agent (LRA; namely PMA or Vorinostat) was followed by collect of supernatant for vRNA quantification (n=5) and co-culture with SupT-1 cells (n=9) in replicate wells (>14 wells). B) vRNA-gag copies/ml of culturing media in presence vs absence of galunisertib (1uM). C) The IUPM calculated based on the frequency of p24+ wells in each condition are shown (pink squares represent PBMC from NIH, triangles represent PBMC from Northwestern, RADAR). Conditions were compared by a 2-way ANOVA followed by the Sidak multiple comparisons test (Significant *p*-values of α<0.05 (*), α<0.01 (**) and α <0.001 (***) are indicated).

To control that addition of galunisertib would not directly impact HIV-1 replication in SupT1 cells or PBMC increasing the chances to detect infection, we performed *ex vivo* infection of SupT1 cells and PBMC from healthy donors in presence of different concentration of galunisertib (Fig S4). Galunisertib had no impact on the HIV-1 growth curves in these *in vitro* infections.

### Blocking TGF-β1 leads to SIV reactivation *in vivo*

To test whether galunisertib might facilitate spontaneous episodes of HIV latency reversal, especially in tissues rich in TGF-β such as the gut, we administered galunisertib at 5mg/kg or 10mg/kg via oral gavage to 7 SIV infected, ART-treated macaques (Fig 3A and Table 2). Macaques were infected intrarectally (2000TCID50), or intravenously (IV) (300TCID50) of SIVmac251 or SIVmac239 (Table 2). All macaques but one were ARV suppressed (plasma viral load; pVL< 100copies/ml) for at least 100 days before galunisertib treatment. Galunisertib was used at 5-10mg/kg, which is equivalent to the dose administered in clinical trials for cancer treatment(*59, 77, 78*) and predicted to be within the therapeutic window in humans by extensive PK/PD studies(*58, 79, 80*). PK in rhesus macaques was evaluated in a separate group of naïve animals (Table 2) and found to be substantially lower than in humans (Fig S5A and (*80*)). However, examination of pSMAD2/3 over total SMAD in PBMC suggested that galunisertib was active in the macaques at these concentrations (∼25% reduction in pSMAD2/3 Fig S5B).

**Table 2.** List of macaques used in the treatment and PK studies.

| *Monkey* | *Sex* | *Weight\**<br>(Kg) | *Age\**<br>(yrs) | *Inoculum* | *Exposure* | *Peak pVL*<br>(copie/ml) | *ART*<br>*Start*<br>(week) | *Weeks*<br>*on*<br>*ART\*\** | *Gal*<br>*Dose%*<br>(mg/kg) | *Treatment*<br>*Length*<br>(days) | *pVL blip on*<br>*ART#*<br>(copies/ml) | *FNA*<br>*before*<br>*Gal* | *FNA*<br>*after*<br>*Gal* |
| --- | --- | --- | --- | --- | --- | --- | --- | --- | --- | --- | --- | --- | --- |
| A14X004 | M | 7.0 | 7 | SIVmac251 | Intrarectal | 1.11x10<sup>5</sup> | 13 | 27 | 5 | 14 | No | Ax | L Ax |
| A14X005 | M | 7.0 | 7 | SIVmac251 | Intrarectal | 4.08x10<sup>4</sup> | 17 | 27 | 10 | 14 | No | Ax | R Ax |
| A14X013 | M | 9.0 | 7 | SIVmac251 | Intrarectal | 1.66x10<sup>5</sup> | 13 | 27 | 10 | 14 | No | L Ax | L Ax |
| A14X027 | M | 7.2 | 6 | SIVmac239 | IV | 3.17x10<sup>5</sup> | 12 | 26 | 5 | 7 | No | L Ax | R ING |
| A14X037 | M | 7.1 | 6 | SIVmac239 | IV | 3.79x10<sup>5</sup> | 12 | 26 | 5 | 7 | No | R Ax | R Ax |
| A14X060 | M | 7.5 | 6 | SIVmac239 | Intrarectal | 1.05x10<sup>6</sup> | 12 | 26 | 5 | 7 | 8.18x10<sup>2</sup> | R Ax | R ING |
| A14X064 | M | 8.0 | 6 | SIVmac239 | Intrarectal | 2.86x10<sup>5</sup> | 12 | 26 | 5 | 7 | 3.06x10<sup>2</sup> | R ING | R ING |
| **PK Study** |  |  |  |  |  |  |  |  |  |  |  |  |  |
| *P83* | F | 9.1 | 14 | N/A | N/A | N/A | N/A | N/A | 2.5 | 1 | N/A |  |  |
| *08M134* | F | 13.5 | 12 | N/A | N/A | N/A | N/A | N/A | 2.5 | 1 | N/A |  |  |
| *08M156* | F | 6.1 | 12 | N/A | N/A | N/A | N/A | N/A | 5 | 1 | N/A |  |  |
| *A6X003* | F | 7.8 | 14 | N/A | N/A | N/A | N/A | N/A | 5 | 1 | N/A |  |  |
| *A8L014* | F | 7.7 | 12 | N/A | N/A | N/A | N/A | N/A | 2.5 | 9 | N/A |  |  |
| *A8L057* | F | 8.5 | 12 | N/A | N/A | N/A | N/A | N/A | 2.5 | 9 | N/A |  |  |
| *A8R095* | F | 7 | 12 | N/A | N/A | N/A | N/A | N/A | 5 | 9 | N/A |  |  |
| *A8T010* | F | 5.0 | 12 | N/A | N/A | N/A | N/A | N/A | 5 | 9 | N/A |  |  |

\*At the beginning of galunisertib treatment
\*\*Before galunisertib
% Twice/day but for P83, 08M134, 08M156, A6X003

**Figure 3.**
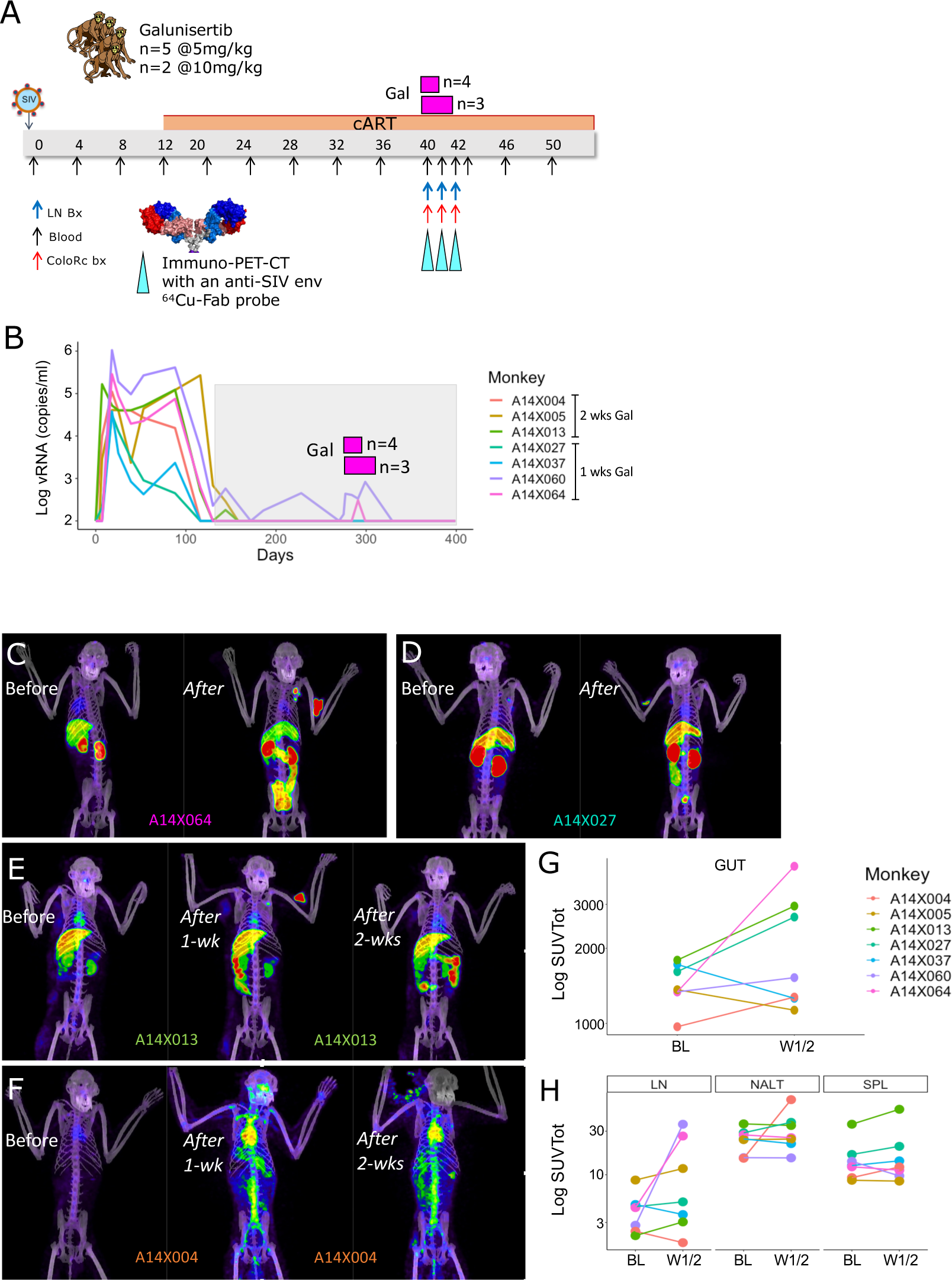
Blocking TGF-β1 leads to HIV-1 reactivation *in vivo*. A) Schematic representation of the macaque studies. SIVmac251 and SIVmac239 infected macaques (n=7) were treated with ARVs starting week 12 post infection. After 26-27 weeks of ART, animals were treated twice daily with 5 or 10mg/kg of galunisertib orally. Colorectal biopsies, fine needle aspirates (FNA) and PET/CT scans with the anti-SIV-env probe ^64^Cu-7D3 were performed before and at the end of the 1 or 2 weeks treatment. The animals that were treated for 2 weeks underwent a 2^nd^ scan at the end of the 1^st^ week of treatment. B) Plasma viral loads measured at NIRC (LLOQ = 100copies/ml) C-D-E-F) Representative images from the PET/CT scans of 4 out of 5 animals with increased PET signal following galunisertib treatment. G-H) SUVtot for different anatomical areas (ROI) are shown before and after galunisertib treatment (Gut includes small and large intestine, LN= axillary LNs, NALT = nasal associated lymphoid tissues, SPL = spleen).

Galunisertib was administered orally twice per day for 1 week for 4 macaques and 2 weeks for 3 macaques. An increase in pVL over 100copies/ml was detected in 2 of the 7 macaques (Fig 3B and Table 2). Additional small changes in pVL occurred below 100copies/ml in the other macaques when pVL was analyzed by ultrasensitive (LLOD 5 copies/ml) assay at Leidos. However, most of the macaques had detectable pVL at baseline when using the ultrasensitive assay and extensive baseline analysis was not performed to determine whether these changes represented a true deviation from baseline.

ImmunoPET/CT images with the validated ^64^Cu-DOTA-F(ab’)_2_ p7D3 anti SIV-env probe(*81, 82*), colorectal biopsies (bx), lymph node (LN) fine needle aspirates (FNA), and blood samples were collected at baseline (day -1 of treatment) and on day 7 or 14 of treatment. Notably, a clear extensive viral reactivation occurred in the gut of 3 of the 7 macaques as documented by the notable increase in PET signal (A14X060, A14X027 and A14X013, Fig 3C-D-E and Supp mov 1, 4 and 5). A smaller increase in the gut was also present in A14X004 and A14X060 (Fig 3G). Viral reactivation also occurred in the LNs of A14X064, A14X013 and A14X060 (Fig 3C and E and Supp mov 3) and NALT, spinal cord and hearth (A14X004, Fig 3F). These increases were captured by the change in total standard uptake values (SUV) in the different anatomical areas as summarized in Fig 3G and H. Interestingly, we observed an increases in the levels of TGF-β in blood after galunisertib treatment (Fig S9A). However, reactivation was not linked to baseline concentrations of circulating total TGF-β (Fig S9B).

Viral reactivation in the macaques was confirmed by large (1 to 3 Log) increases in cell associated vRNA (CA-RNA) observed when colorectal biopsies and LN FNA were assayed for RNA (Fig 4A). Notably, the increase in rectal biopsy CA-RNA correlated well (R=0.85; Fig 4B; Upper) with the increase in SUVtot in the gut region. This occurred even though biopsies were collected from the colorectal area while the SUVtot for the gut represent the signal in the entire intestine. In contrast, no correlation was present between vRNA in the FNA that sampled a single lymph node (Table 2) and the SUVtot signal in both axillary LN (Fig 4B; Lower).

**Figure 4.**
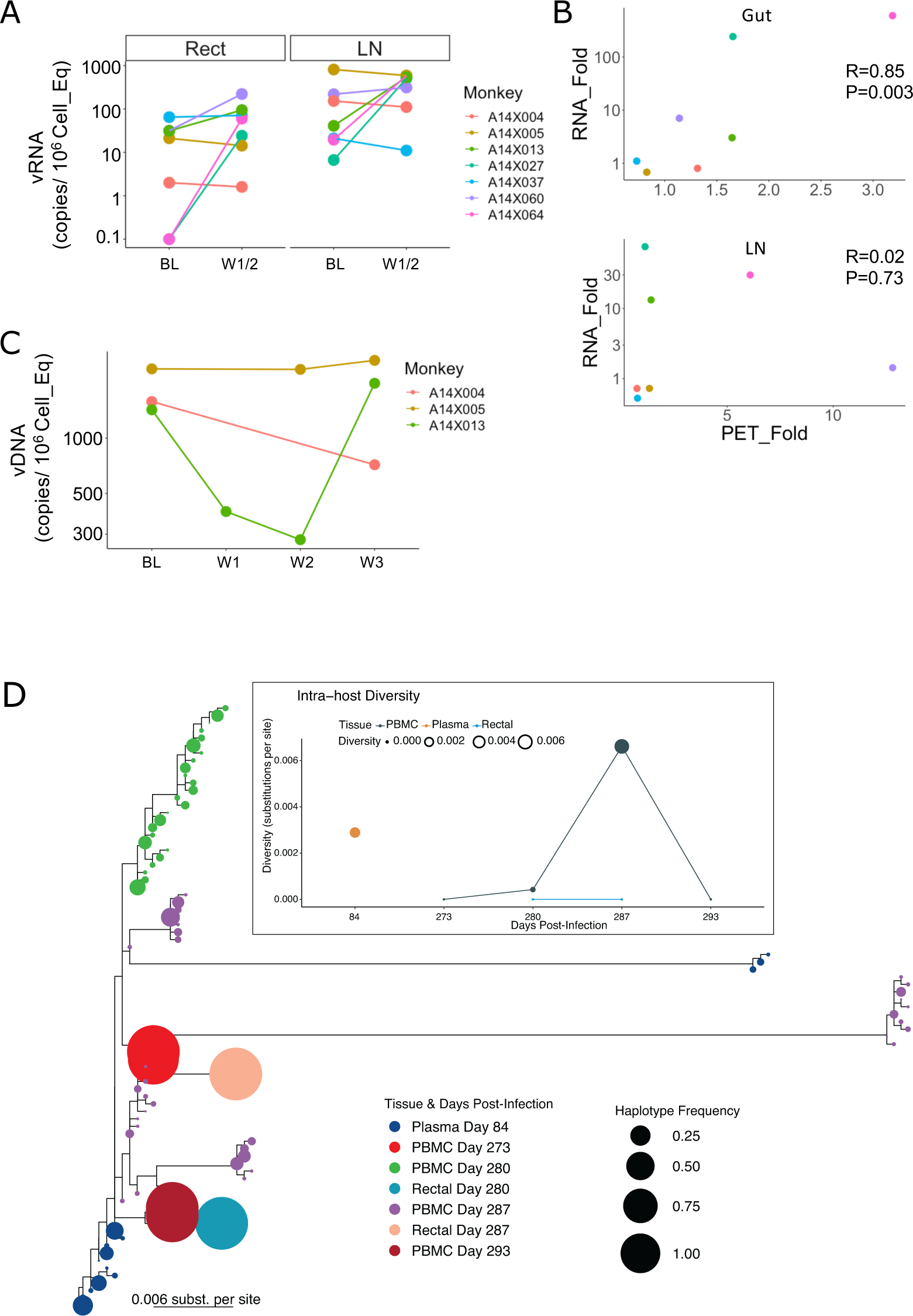
Increased PET-signal corresponds to increased vRNA. A) Copies of cell-associated unspliced vRNA normalized on 10^6 cells diploid genome equivalent are shown for colorectal biopsies and LNA before and after galunisertib treatment. B) The correlation between the fold increase in SUVtot for the gut (above) and LN (below) and the fold increase in vRNA copies in rectal biopsies and FNA (only 1 of the axillary LNs was sampled) are shown. Pearson’s correlation coefficient and p values are indicated in each graph. C) Copies of vDNA per 10^6^ cells-equivalent are shown for the PBMC of the 3 animals that were treated with galunisertib for 2 weeks. W3 time point represents a sample collected 7 days after the last galunisertib administration. D) Intra-host viral quasispecies evolution before and after galunisertib treatment in A14X013 is shown. Inferred ML tree with the reconstructed the intra-host quasispecies in PBMCs, rectal tissues, and blood samples using deep sequencing of the gag gene. Circles at the tips represent each haplotype and are colored by time post-infection and sample type (D84 is plasma right before ART initiation; D273 is PBMC on the day of first galunisertib treatment, collected before treatment; D280-287-290 represent 7, 14 and 20 days after galuniertib initiation). Circle size indicates the frequency of the haplotype in the quasispecies. **Insert** represents the spatio-temporal evolution of the viral population diversity at each compartment and time point. Both y-axis and circle size indicate diversity of the quasispecies and data from the same sample type are connected by lines. Intra-host viral diversity was calculated as the weighted average of pairwise distances between every haplotype weighted by their frequency in the population in substitutions per site.

In some macaques vRNA increase was paralleled by a small increase in CA-DNA (Fig S7 A; eg in A14X060 for colorectal bx and A14X027 and A14X064 for LN). In other macaques CA-RNA increase was not accompanied by an increase in CA-DNA and CA-DNA varied substantially between baseline and post-galunisertib treatment. However, the ratio CA-RNA/DNA increased substantially in at least 2 macaques (A14X027 and A14X064, Fig S7B).

We examined the levels of CA-vDNA in the PBMC of the 3 macaques that were treated for 2 weeks (samples were not available for the animals treated for 1 week). Interestingly, we noted a drop in the CA-vDNA levels in the PBMC of A14X013. Since this macaque had a strong viral reactivation signal via PET (Fig 3E), we considered this drop interesting and worth further evaluation by viral sequencing. The increase in CA-vRNA in the tissues of this macaque was balanced by an increase in CA-DNA, suggesting that galunisertib may have induced recirculation of infected cells from blood to tissues and vice versa. This was supported by the sequencing data that demonstrated a substantial increase in the diversification of the intra-host viral quasispecies obtained from A14X013 PBMCs during the two weeks of galunisertib treatment (Fig. 4D). The inferred phylogeny of the intra-host quasispecies at the different time points and anatomical locations suggest the expansion of multiple viral lineages at week 1 and 2 after galunisertib initiation in PBMC (Fig 4D). Viral diversity was low at the time of galunisertib initiation in PBMC. This was followed by the high levels of diversification in blood while observing compartmentalization between PBMC and rectal tissues that additionally displayed low diversity populations. At week 3 after galunisertib treatment initiation (1 week after the end of the treatment), viral diversity in blood dropped and the new dominant viral populations in blood were closely related to the later viral populations in rectal tissues detected a week earlier. Overall, this suggests a significant viral expansion during galunisertib treatment of multiple infection foci and migration of viral populations between tissue and blood in A14X013.

### Blocking TGF-β1 enhances SIV-specific responses

Since TGF-β is primarily an immunosuppressive factor, blocking TGF-β1 signaling was expected to impact phenotype and function of immune cells independent of SIV infection. Indeed, a preliminary mRNAseq analysis of CD4^+^ T cells from before and after (6hrs) Galunisertib treatment in 3 naïve macaques suggested notable changes in pathways regulating immune functions such as leukocyte activation and adaptive immune system (downregulated) and regulation of DNA binding, MAPK, and FAK pathways and TCR signaling (upregulated) (Fig S8). Hence, we investigated the impact of TGF-β1 blockade with galunisertib on SIV-specific T cell and B cell responses. T cell responses were probed via classical PBMC stimulation with gag and env peptide antigen followed by detection of intracellular INF-γ, TNF-α, and IL-2 by flow cytometry. We found a significant increase in the frequency of INF-γ producing CD4^+^ and CD8^+^ T cells (Fig 5A) following both gag and env peptides stimulation and in TNF-α producing CD4^+^ T cells following Env stimulation (Fig 5A). However, we observed no increase in the frequency of cells producing multiple cytokines (pluripotent cells). Subset analysis using a CD95 memory marker was not performed.

**Figure 5.**
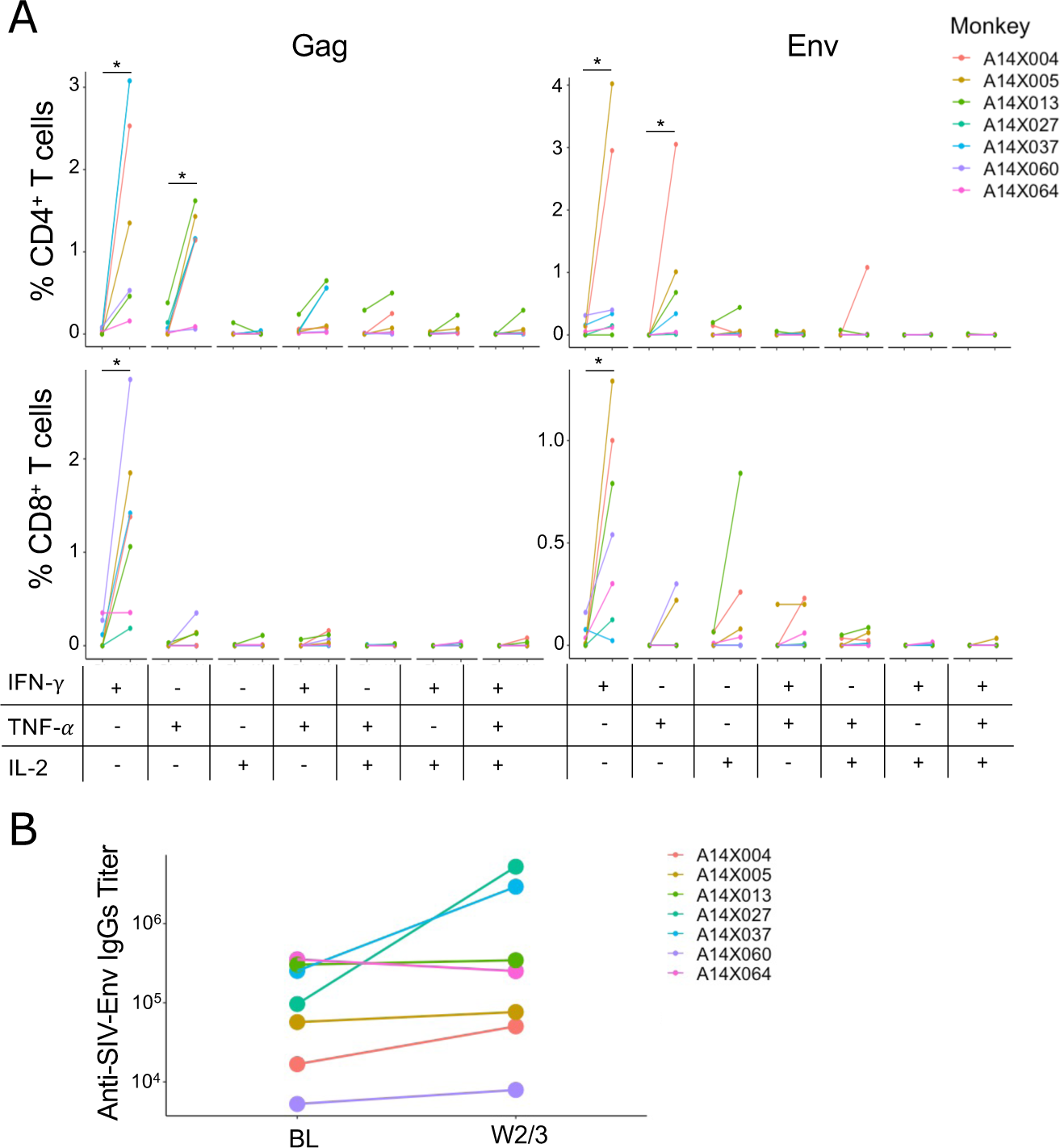
Blocking TGF-β1 stimulates SIV-specific responses. A) The frequency of CD4^+^ T cells and CD8^+^ T cells that express the cytokines indicated below the graphs in response to stimulation with a pool of 15-mer overlapping peptides from SIVmac239 gag and env are shown for before and after galunisertib treatment. The frequency of each population was calculated after subtraction of a DMSO unstimulated control performed in parallel with each sample. Wilcoxon matched pairs signed rank test was used to compare data before and after galunisertib. B) SIV-env titers in plasma before and after galunisertib treatment are shown for each animal (Significant *p*-values of α<0.05 (*), α<0.01 (**) and α <0.001 (***) are indicated).

Finally, in 3 of the 7 macaques there was a notable (0.5 to 1 Log) increase in the titer of SIV-env specific antibodies following galunisertib treatment (Fig 5B). However, this increase did not occur in macaques with large viral reactivation as measured by TotSUV increase. Rather antibody titers increased in animals with no change (A14X037) or relatively lower increase (A14X027 and A14X004) in gut TotSUV (Fig 3G).

## Discussion

HIV-1 persists in different cell types during ART and it is now clear that the mechanisms of viral persistence are heterogeneous. HIV-1 transcriptional activity varies in different cell types and is dependent, at least in part, on tissue location and environmental cues. HIV-1 latency in T cells is maintained through diverse mechanisms that include blocks in transcriptional elongation, completion, and splicing(*4*). The differential role of these blocks on HIV-1 expression in T cells depends on the tissue and microenvironment where infected cells reside and the same factors may also influence the ability of HIV-1 to emerge from latency (*83, 84*). Similar heterogeneous, tissue/environment-dependent mechanisms may be at play in regulating HIV-1 replication in other HIV cell reservoir types such as myeloid cells. Finally, the long-term persistence of these infected cells may be dependent on homeostatic(*85, 86*) or antigen-driven proliferation(*5, 87-89*) with a potential contribution of promoter insertion mutagenesis(*90, 91*). The different cellular processes that regulate the dynamics of HIV-1 reservoir remain to be fully clarified. However, a common characteristic shared by HIV-1 target cells of both T and myeloid cell lineages that constitute the HIV-1 “invisible” reservoir is their “resting” phenotype. TGF-β represents a common factor released at high level in PLWH. It limits the ability of HIV-1 infected cells to respond to activation stimuli and raises the threshold for TCR-driven proliferation,(*48*) while not fully impairing homeostatic proliferation(*92*). The capacity of TGF-βto broadly inhibit cell activation in several different potential HIV-1 cellular reservoirs, while not fully blocking the ability of infected cells to proliferate led us to test the hypothesis that TGF-β might be a major factor that contributes to the low inducibility of intact proviral sequences in a wide range of cell types(*20*). We reasoned that inhibiting TGF-β signaling might decrease the threshold for transcription and translation of HIV-1 proteins in cells activated by different stimuli both *ex vivo* (in presence of exogenous TGF-β1, including the TGF-β1 present in FBS(*30*)) and *in vivo*. In tissues, especially mucosal tissues such as the gut, TGF-β is a major player and regulator of tissue homeostasis(*93, 94*). Moreover, the gut is a site of constant antigenic exposure. This is especially true in HIV infected subjects where compromised gut barrier function is a source of inflammatory stimuli that likely drives stochastic HIV-1 reactivation. Indeed, *in vitro*, using the classical cell line models ACH-2 and U1, we demonstrated that addition of exogenous TGF-β decreases the ability of PMA, a powerful PKC activator, to induce HIV reactivation and, as consequence, fewer cells expressed HIV antigens in presence of TGF-β. This effect was blunted when TGF-β was added in presence of galunisertib. In some cases, we detected enhanced HIV-1 reactivation in the presence of galunisertib alone, which likely reflected galunisertib-mediated inhibition of FBS-derived TGF-β.

We also demonstrated that TGF-β acts both directly and indirectly on HIV-1 infected cells. Indeed, TGF-β-treated moDCs exhibited a reduced ability to reactivate HIV-1 from latently infected T cells. Historically, PBMCs and CD4^+^ T cells have been cultured in the presence of FBS. Both FBS-derived TGF-β and TGF-β released by HIV-infected PBMC likely exert an inhibitory effect on HIV reactivation *ex vivo* in qVOA-type assays. This explains the activity of galunisertib in increasing the frequency of latency reactivation in our qVOA. Importantly, our qVOA was a highly simplified and modified version of the classical qVOA(*76*). A major limitation of this work is that we have only tested PMA and Vorinostat as LRAs. We did not determine whether inhibiting TGF-β signaling may also increase HIV reactivation following treatment with BRD4 inhibitors, HMT inhibitors or TLR inhibitors. TGF-β1 is known to inhibit TLR-signaling, so it is likely that galunisertib will enhance the LRA activity of TLR agonists. Its impact on other LRA classes is less clear, and will need to be addressed in future studies.

The most intriguing result that we obtained involved the ability of galunisertib to drive SIV reactivation *in vivo* in aviremic, SIV infected macaques. While we detected a clear SIV env signal in tissues by PET/CT following galunisertib treatment in 5 out of 7 animals, we detected an increase in VL in only 2 out of the 7 animals. A14X060, whose viremia had not been continuously suppressed in blood before galunisertib treatment, had a very high (>2 Log) increase in pVL with a relatively low increase in gut SUVtot. However, this animal had a substantial increase in PET signal in the LNs which may explain the increase in pVL. In contrast, in the other 4 animals that showed reactivation in tissue there was minimal or no increase in vRNA in blood. Considering that we did not sample blood every day, rather every 3-4 days, it is possible that we missed a transient increase in pVL. However, it is also possible that viral expression/reactivation was localized in tissues and not enough virus reached the blood to be detected. Importantly, the observed increase in SUVtot in the gut was reflected in the increase in vRNA in colorectal biopsies. This occurred despite the highly focalized sampling of biopsies compared with measuring the overall PET signal in the entire organ, warranting further study and confirmation. However, it is important that the data collected with the two different techniques agree with each other. In contrast, the lack of correlation between increase in PET signal in lymph nodes and the increase in vRNA in FNA may be due to the FNA technique involving sampling a single site, not necessarily axillary, while the PET signal that we could isolate with the MIM software encompassed only the axillary LNs.

Some insights into the dynamics of viral rebound on ART during galunisertib treatment comes from the sequencing done on A14X013. This animal had a large increase in gut SUVtot detected by immunoPET, but no increase in pVL was detected with our sampling schedule. Yet we found evidence of increased viral diversity during galunisertib treatment particularly in blood that concluded with a new dominant viral population closely related to those detected in rectal tissue. Although this analysis was done only in 1 macaque, it suggests that galunisertib may induce viral expansion of multiple infection foci and migration of viral populations between tissue and blood.

Finally, we observed an increase in SIV-specific immune responses that appeared to be driven by the galunisertib treatment. Since these increases were also detected in the animals that exhibited no detectable rebound (A14X005 and A14X037), they may not have been directly related to increases in viral antigen. This is intriguing and suggests potential direct effect of inhibiting

TGFBR1 signaling on SIV-specific CD4 and CD8 cells. An increase in T cell function was also detected during the preclinical and clinical development of galunisertib(*57, 58*). Whether this increased immune response could translate into increased purging of the viral reservoir was not the focus of this first proof-of-concept study and remains to be determined. In any case, the combination of viral reactivation and enhanced cellular and humoral responses in this first pilot study is highly promising and suggests that galunisertib may have an impact on the viral reservoir alone or in combination with other strategies to enhance immune functions killing virus-expressing cells.

The studies presented herein have several limitations. They include the use of cell lines as models for HIV latency, which may not fully recapitulate the mechanisms of latency *in vivo*. Moreover, as mentioned above only PMA and SAHA were tested in combination with galunisertib in a limited number of PBMC samples from aviremic PLWH. Additional studies will be needed to dissect potential differences in the activity of galunisertib on different mechanisms of HIV-1 reactivation. Regarding the *in vivo* studies, the limited number of animals and lack of prolonged follow-up or ART interruption as well as relatively short treatment course are limitations that can be addressed in future studies now that we have determined that galunisertib is able to mediate virus reactivation. Our dosing was based on pre-clinical studies in other animals and on the therapeutic index used for clinical studies in humans. However, the concentration of galunisertib in the blood of macaques is at least 10-fold lower than the average concentration detected in human plasma after similar dosing. Future work will require more detailed PK/PD studies in macaques. Another drawback of our study is that we are missing analysis of cell phenotypes in PBMC and tissues before and after galunisertib administration *in vivo*. This was not done because of limited sample availability. In future studies, it will be important to determine whether galunisertib treatment leads to an increase in markers of immune activation or an increase in the expression of inflammatory markers. It is noteworthy that viral reactivation did not correlate with higher baseline TGF-β concentrations in blood. Hence, the effect of galunisertib on the virus appears to be independent of TGF-β levels in plasma and the levels in plasma may not directly reflect the levels in tissues. Moreover, we monitored total TGF-β and the levels of active TGF-β may be different. Future studies should include monitoring of TGF-β in tissue at baseline to understand whether galunisertib may work better in the context of high baseline tissue TGF-β levels.

In conclusion, we have demonstrated that TGF-βinhibits HIV reactivation from latently infected cells, an activity that may impact *ex vivo* assays aimed at quantifying the size and dynamics of reservoirs. We further demonstrate that a small molecule TGFBR1 inhibitor, galunisertib, in advanced stages of clinical development, can promote viral reactivation *ex vivo* and *in vivo*. Measurements of pVL did not detect reactivation *in vivo* in all instances, likely due to insufficient sensitivity. Our work demonstrates that immunoPET/CT imaging can provide important additional information regarding the efficacy of HIV-1 cure strategies. Because this was a first attempt at using an antagonist of TGF-β to reactivate the HIV-1 reservoir *in vivo*, we expect that optimization of treatment protocols will lead to improved activity. Nonetheless, our results are both surprising and promising and may lead to novel therapeutic interventions to reduce or eliminate the HIV-1 reservoir and bring the field closer to a cure.

## Methods

### Study design

A total of 15 adult male and female Indian *Rhesus* macaques (*Macaca mulatta*; Mamu A*01, B*08 and B*17 negative) were used for the *in vivo* studies described (Table 2). All rhesus macaques used in these studies were selected form the colonies bred and raised at the New Iberia Research Center, University of Louisiana at Lafayette. Macaques were housed and maintained at NIRC in accordance with the rules and regulations of the Committee on the Care and Use of Laboratory Animal Resources. Treatment of all macaques were in accordance with NIRC standards and any abnormal observations and/or signs of illness or distress were reported per NIRC SOPs. All animals were negative for SIV, simian T cell leukemia virus and simian retrovirus, previous to the study. All animal experiments were conducted following guidelines established by the Animal Welfare Act and the NIH for housing and care of laboratory animals and performed in accordance with institutional regulations after review and approval by the Institutional Animal Care and Usage Committees (IACUC) of the University of Louisiana at Lafayette (2016-8761-067; protocols 8761-1709 and 8761-1802). All efforts were made to minimize suffering and stress. Appropriate procedures were performed to ensure that potential distress, pain, discomfort, and/or injury were limited to that unavoidable in the conduct of the research plan. Ketamine (10 mg/kg) and/or telazol (4 mg/kg) were used for sample administration, collection of samples and conducting image collections. Analgesics were used when determined appropriate by veterinary medical staff. Monkeys were fed monkey chow supplemented with fresh fruit or vegetables and water ad libitum. Rhesus macaques (n=7) were infected with SIVmac251 or SIVmac239 rectally or intravenously (Table 2) and ART (Tenofovir [PMPA], Emtricitabine [FTC] and Dolutegravir [DTG]) was initiated between week 12 and 17 post infection. Galunisertib treatment was initiated after 26 or 27 weeks on ART for 7 or 14 days. Galunisertib powder (MedChemExpress – MCE, NJ, USA) was dissolved in water and given by oral gavage twice daily at 5mg/kg or 10mg/kg (Table 2). Blood viral load was monitored biweekly before and during cART and every 3-4 days during Galunisertib treatment. Colorectal biopsies and LN FNA were collected before and after galunisertib treatment.

PET/CT imaging with an 64Cu-DOTA-F(ab’)2 p7D3 anti SIV-env probe were performed before Galunisertib treatment, after 7 days in all macaques and after 14 days in the macaques that were treated for 14 days. Briefly, a solution of 64Cu was incubated with DOTA-F(ab’)2 p7D3 antibody reconstituted with 50uL of 1M NH4OAc pH of 5.5 and incubated for 1 hour at 37C; then purified with a Zeba desalting spin column (10K MWCO, Thermo Fisher) and finally eluted with sterile saline prior to injection. The labeled probe was injected via IV, 24h before PET-CT imaging. For taking PET/CT scans, the macaques were sedated with ketamine HCL (10mg/kg), and/or telazol (4-8mg/kg) via intramuscular and supplemented with ketamine (5mg/Kg) as needed to perform the procedure. The macaque’s body was immobilized in dorsal recumbency on the scanner table and PET/CT scans were acquired using a Philips Gemini TF64 PET/CT scanner. The final CT image was compiled from 250 to 300 slices, depending on macaque size.

PET Image analysis was performed using the MIM software. PETCT fusions were generated scaled according to calculated Standardized Uptake Values. The SUV scale for the PET scans was selected based on the overall signal intensity of the PET scans (whole body), and the CT scale was selected for optimal visibility of the tissues. All images and maximum image projections were set to the same scale for visual comparisons.

Regions of interest (ROI) were isolated using a combination of the Region Grow function and manual contouring on a representative scan. These regions were the copied onto subsequent scans of the same animal using a specialized developed workflow. This workflow uses the CT scans to map the selected ROI and locate that corresponding volume in subsequent scans. Manual adjustments were then used to counter any changes in the animals’ orientation between scans. The signals of the whole body as well as within these ROI were quantified, and Max, Mean, and Total SUVs were calculated for further analyses. Organs of high and unwanted signal (kidneys and liver) were masked using the Region Grow function for selecting the boundaries of the organ signal and the Mask function to reduce the signal within the contour to background levels (for A14X004 only).

### SIVmac251 and SIVmac239 preparation and administration

Viral stocks were propagated and titrated in Rhesus PBMC and prepared for either intrarectal or intravenous dosing in a biosafety cabinet and diluted in medium. Rhesus macaques were inoculated intravenously with 300 TCID50 of SIVmac239 or intrarectally with 2000 TCID50 of SIVmac251 or SIVmac239. Intravenous injections or intrarectal administration were done using 1mL of diluted virus and macaques were under sedation.

### In vitro models of HIV latency reactivation

ACH2 cells and U1 cells were obtained through the NIH HIV Reagent Program, Division of AIDS, NIAID, NIH (contributed by Dr. Thomas Folks). Cells (10^5^/well) were cultured in triplicates in RPMI with 10% FBS and Pen/Strep in presence of TGF-β1 (10ng/ml R&D systems), TGF-β1 and galunisertib (1μM MedChemExpress), galunisertib alone or mock treated. In the HIV latency reactivation conditions, PMA (Phorbol 12-myristate 13-acetate, SigmaAldrich) was added at 100ng/ml together with galunisertib or TGF-β1. After 18hrs, cells were stained intracellularly for p24 using the KC57 antibody (NIH HIV Reagent Program) as previously described (*95*).

Blood from de-identified donors was obtained from the New York Blood Bank and used as source of CD4^+^ T cells for the primary model of HIV latency *in vitro*. Total CD4^+^ T cells were isolated with the CD4^+^ T cells isolation kit (Miltenyi) and activated with 5μg/ml of PHA and 50U/ml of IL-2 for 3 days. After washing out the PHA, cells were infected with 200TCID50 of HIV-ADA (NIH HIV Reagent Program) in 12 or 24 wells/plates and spinoculated at 1200g for 2hrs at 25°C. After additional 3hrs of incubation at 37°C, viral inoculum was washed and cells plated in presence of T20 (Enfurtivide acetate salt, 1μM NIH AIDS reagent program). After 48hrs, cells were treated with PMA (100ng/ml) in presence and absence of galunisertib (1μM) overnight and stained for p24 intracellularly as described above.

CD14+ cells were isolated from PBMC using the Miltenyi CD14+ beads. Monocyte-derived DCs (moDCs) were obtained from CD14+ cells cultured in GM-CSF and IL-4 as previously described(*73*). At day 4 of moDCs differentiation, TGF-β1 (10ng/ml R&D systems) was added to the moDCs to generate the TGF-β-DCs (or both TGF-β1 and galunisertib). At day 6, 2 days after TGF-β1 addition, moDCs were phenotyped (in 3 donors; See Supplemental Table 1 for antibodies list) and co-cultured with eFluor670 (eBioscience) -labelled U1 at 3:1 U1:DC ratio (5 donors). After 18hrs of co-culture, cells were stained for p24 intracellularly as described above.

### Quantitative viral outgrowth assay

The standard qVOA assay(*76*) was extensively modified. PBMC from aviremic donors were used in absence of resting CD4+ T cells isolation. De-identified PBMC were obtained either from NIAID protocol 14-I-0039 or 10-I-N048 or from the Northwestern University RADAR cohort.

PBMC were thawed and cultured in RPMI 10% FBS at 3 × 10^6^/ml. Cells were activated with PMA (100ng/ml or 10ng/ml) or Vorinostat (1μM, NIH HIV Reagent Program) in presence of T20 (HIV Reagent Program) and IL-2 (100U/ml) and galunisertib (1μM) or mock (DMSO). After 18hrs, cells were washed twice in PBS and plated at 3×10^5^ / well (>14 replicate wells; a number where we expected <1 HIV latently infected cell/well(*96, 97*)) in a flat-bottom 96 wells/plate. 10^5^ SupT1-R5 (M10 clone, HIV Reagent Program) were added in each well and galunisertib (1μM) or an equal volume of DMSO were added to the respective wells in each condition. 50μL of supernatant was collected after 7, 14 and 21 days of co-culture and monitored for p24 presence by p24 ELISA (PerkinElmer). IUPM were calculated based on the number of positive wells using the IUPM calculator provided by Dr. Siliciano(*98*). Supernatant from stimulated PBMC before co-culture with SupT1 cells was tested by *gag*-RT-qPCR(*99*).

For control experiments of the impact of galunisertib on HIV infection of SupT1 cells and PBMC, SupT1 cells and PHA-activated PBMC were cultured respectively at 1 and 2millions/ml in R10 (in presence of 20U/ml of IL-2 for the PBMC) and infected with 50 TCID50/10^6^ cells of HIV-Bal (0.01 MOI). HIV replication was monitored by p24 ELISA.

### Plasma and Tissue SIV Viral loads

Blood was collected in EDTA tubes and plasma was separated by density gradient centrifugation and used for the determination of plasma VL by SIVgag qRT-PCR at NIRC or at Leidos (Quantitative Molecular Diagnostics Core, AIDS and Cancer Virus Program Frederick National Laboratory). Tissue VL from snap frozen colorectal biopsies and LN FNA were performed at Leidos (A14X004, A14X005 and A14X013) as described in(*100*) or, for the remaining macaques and from PBMC of all macaques they were performed as described in (*99*).

8 naïve macaques (Table 2) were used to collect plasma and PBMC at different time points after galunisertib treatment. 4 animals received galunisertib (2.5mg/kg, n=2 or 5mg/kg, n=2) only once and blood was collected at 1hr, 3hr and 6hrs, 24hrs and 48hrs. 4 animals received galunisertib (2.5mg/kg, n=2 or 5mg/kg, n=2) twice daily for 9 days and samples were collected 1hr, 3hr and 8hrs after administration on day 1 and on day 9 right before and 1hr and 3 and 6hrs after the last administration. Plasma concentrations of galunisertib were measured by the Northwestern IMSERC (Integrated Molecular Structure Education and Research Center) utilizing liquid chromatography mass spectrometry (Shimadzu Kyoto, Japan ultra-high performance liquid chromatography – UHPLC – system and SCIEX Framingham, MA Qtrap 6500+ triple quadrupole mass spectrometer). All standard, quality controls (QC) and experimental samples were prepared and ran in duplicate using 20 μL of plasma (spiked with appropriate concentration of galunisertib if a standard or QC) adding 15 μL of 8% phosphoric acid solution, 200 μL of acetonitrile (to crash plasma proteins) spiked with tramadol as an internal standard, vortexed for 1 minute, and centrifuged for two minutes at 13,200 rpm before transferring the supernatant to a 96 well plate for LCMS analysis. The standard curve was prepared as a 10-point curve from 1.0 to 1,000 ng/mL with three QCs. For the LCMS method, a 1 μL injection was performed of the resulting supernatant separated on a Phenomenex (Torrance, CA) Kinetex® Biphenyl 2.6 um 100A LC column (50 × 2.1 mm). The separation gradient was performed using 0.1% formic acid in water (A) and 0.1% formic acid in methanol (B) mobile phases following a gradient at 0.6 mL/min flow rate of: 0.5 minutes 10% B, 4.0 minutes 85% B and hold for 1.0 minutes, and returning to starting conditions of 10% B at 5.1 minutes holding for 1.4 minutes. All samples and standards were processed in duplicate with the average area under the curve (AUC) being used for quantitation reporting. After internal standard correction, all samples and standards outside of a 20% CV between preparation replicates were re-aliquoted and rerun in duplicate to meet this requirement. All standards and QCs were required to be reported within a 20% accuracy tolerance following internal standard correction. The resulting limit of quantitation (LOQ) was 1.0 ng/mL.

For PD studies, the levels of phosphorylated SMAD2/3 were measured using the PathScan PhospoSMAD2/3 ELISA kit (Cell Signaling Technology, CST) over the levels of Total SMAD2/3 in cell lysates Total SMAD ELISA Kit (Cell Signaling Technology, CST) normalized on protein content.

### Sequencing of SIV genome in PBMC, rectal biopsies and plasma from A14X013

DNA was extracted from PBMC and rectal biopsies using the DNAeasy blood and tissue kit (Qiagen), while RNA was extracted from plasma using the QIAamp Viral RNA Mini Kit (Qiagen). Extracted RNA was immediately reverse transcribed using SuperScript™ VILO™ cDNA Synthesis Kit (Invitrogen). Q5® Hot Start High-Fidelity 2X Master Mix (NEB) was used to amplify the *gag* gene with primers Outer-Gag-F (5’-GAAGCAGGAAAATCCCTAGCAG-3’) and 8R (5’-CCAACTGACCATCCTTTTCCATCTTT-3’) followed by a second amplification round using primers 1F (5’-TCCTGAGTACGGCTGAGTGAAG-3’) and 7R (5’-TCCTATTCCTCCTACTATTTTTGGGGT-3’)(*101*). PCR cycling conditions were as follows: 98ºC for 30 s, 35 cycles of 15 s at 95ºC, 30 s at 65ºC, 2 min at 72ºC and a final extension of 72ºC for 2 min. Sequencing library preparation of the obtained amplicons was performed using the SeqWell plexWell 384 kit per manufacturer’s instructions. Pooled libraries were sequenced on the Illumina MiSeq using the V2 500 cycle kit. We subsequently performed reference-based mapping assembly of the sequencing reads using the HAPHPIPE protocol (PMID: 33367849) and SIVmac239 reference genome (M33262.1). With this protocol, reads were trimmed to remove adapters and low-quality sequences using Trimmomatic v0.39 (PMID: 24695404) and error-corrected using SPAdes v3.13.1 (PMID: 22506599). PBMC samples were sequenced twice to confirm the results obtained.

### Phylogenetic and Diversity Analysis

With the assembled reads we performed probabilistic inference of intra-host viral quasispecies for each sample obtained from A14X013 using QuasiRecomb (PMID: 23383997). The sequences of the inferred viral haplotypes from each quasispecies were aligned using MAFFT v7.453 software and a Maximum Likelihood (ML) phylogeny was reconstructed with IQ-Tree v2.0.5 to track the spatio-temporal evolution of the intra-host viral populations. The final tree representation was performed with the R package ggtree v3.2.1 (https://doi.org/10.1111/2041-210X.12628). Intra-host diversity for each sample analyzed was calculated using DistanceCalculator in Biopython and pairwise distances weighted by the estimated frequency of the haplotype with an in-house python script.

### Anti-SIV T cell responses

Frozen PBMC collected right before the first galunisertib administration and right after were thawed in AIM V medium (Thermo Fisher) plated on a plate pre-coated with 2.5μg/ml goat anti mouse (GAM) IgGs and cross linked with 10μg/ml anti-CD28 and anti-CD49d antibodies (Sigma Aldrich). Cells were stimulated either with SIVMAC239 GAG peptide pool (2μg/ml) or with with SIVMAC239 ENV peptide pool (2μg/ml AIDS reagents program, Division of AIDS, NIAID, NIH) or the same volume of DMSO as baseline control. Pooled cells stimulated with 1μg/ml PMA/Ionomycin were used a positive activation control. One hour later 10μg/ml BFA and monensin (GolgiStop, BD Biosciences) were added to each well. After 5 hours, cells were transferred to a FACS plate and stained with the LIVE/DEAD Aqua viability dye (Thermo Fisher Scientific) and antibodies against cell surface and cytokines (Supplemental Table 1).

### Anti-envelope antibodies

96-well plates were coated with the SIV gp130 Recombinant protein (AIDS reagents program, Division of AIDS, NIAID, NIH at 1ug/ml in coating carbonate buffer 2 hours at room temperature (RT). After washing the plates 4 times with washing buffer, the plates were incubated in blocking buffer (2% BSA in PBS) overnight. Plates were washed and dilutions of test sera or control sera were added to the plate starting at 1:50. Sera were incubated at 37°C for 30’ and plate washed and detected with anti-Rhesus IgG HRP (NHPRR) 30’ at 37°C. After washing and substrate incubation at RT for 30’ plates were read at 450/650nm.

### Statistics

Data from different conditions of ACH-2 and U1 experiments were compared using the Kruskal-Wallis ANOVA test followed by the Dunn’s test corrected for multiple comparisons. For the QVOA experiments, IUPM with and without galunisertib were compared in the LRA different conditions using 2-way ANOVA followed by the Sidak multiple comparisons test. Before and after galunisertib SUV, tissue VL data as well as anti-envelope titers and frequency of cells producing different cytokines combination following peptide pool stimulation were compared using the Wilcoxon non-parametric matched-pair signed rank test. DC phenotype between TGF-β-DC and moDCs was compared using the ratio paired t-test. Statistical analysis was performed using Prism GraphPad 9.3 and R.

## Acknowledgments

We are very grateful to Jeffrey Lifson, William Boshe, Rebecca Shoemaker and the team of the AIDS and Cancer Virus Program (Leidos Biomedical Research, Inc.) for supporting this work with timely plasma and tissue viral load measurements. We thank the RADAR study team for providing samples. This work made use of the Clinical Pharmacology Core at Northwestern University part of the Integrated Molecular Structure and Education Research Center (IMSERC). We acknowledge the outstanding help and assistance of the professional staff, animal technicians and caretakers of the New Iberia Research Center and the staff of the Robert H. Lurie Comprehensive Cancer Center Flow Cytometry Core Facility of the Northwestern University for their assistance with flow cytometry analysis. We thank Dr. Richard D’Aquila, Professor of Medicine at Northwestern University for editing the manuscript.

## Funding Statement

This research was funded in part by grants NIH-R01 AI139288 and R01 AI111907 to FV and PJS, R56 AI157822 to EM, P01 AI131346 and P30 AI117943 to TJH. The work of IS, JA, and CC was supported by the intramural research program of NIAID/NIH. The Clinical Pharmacology Core at Northwestern University (IMSERC) received support from the NIH (1S10OD012016-01 / 1S10RR019071-01A1), Soft and Hybrid Nanotechnology Experimental (SHyNE) Resource (NSF ECCS-1542205), the State of Illinois, and the International Institute for Nanotechnology (IIN).

## Authors Contribution

SS MSA JMA IF MM CTT MRA EM performed the experiment and sample assays and analyzed the data; YST and PS analyzed the PET/CT images; BO and AG performed the galunisertib PK analysis; AM, CMC, MLS, DB an FJV coordinated the macaque studies, collected samples and performed PET/CT imaging; LS, JFH and RLR performed the viral sequencing and RLR analyzed the data; JA, CC, IS contributed to the *ex vivo* studies, data interpretation and draft manuscript; TJH, FJV and EM interpreted the data; EM conceptualized the studies and wrote the manuscript.

## Disclaimer

Authors declare no conflict of interest.

## Notes

### Competing Interest Statement

The authors have declared no competing interest.

