## Supplemental Figure for "Blockade of TGF-β signaling reactivates HIV-1/SIV reservoirs and immune responses *in vivo*"

### Slide 1
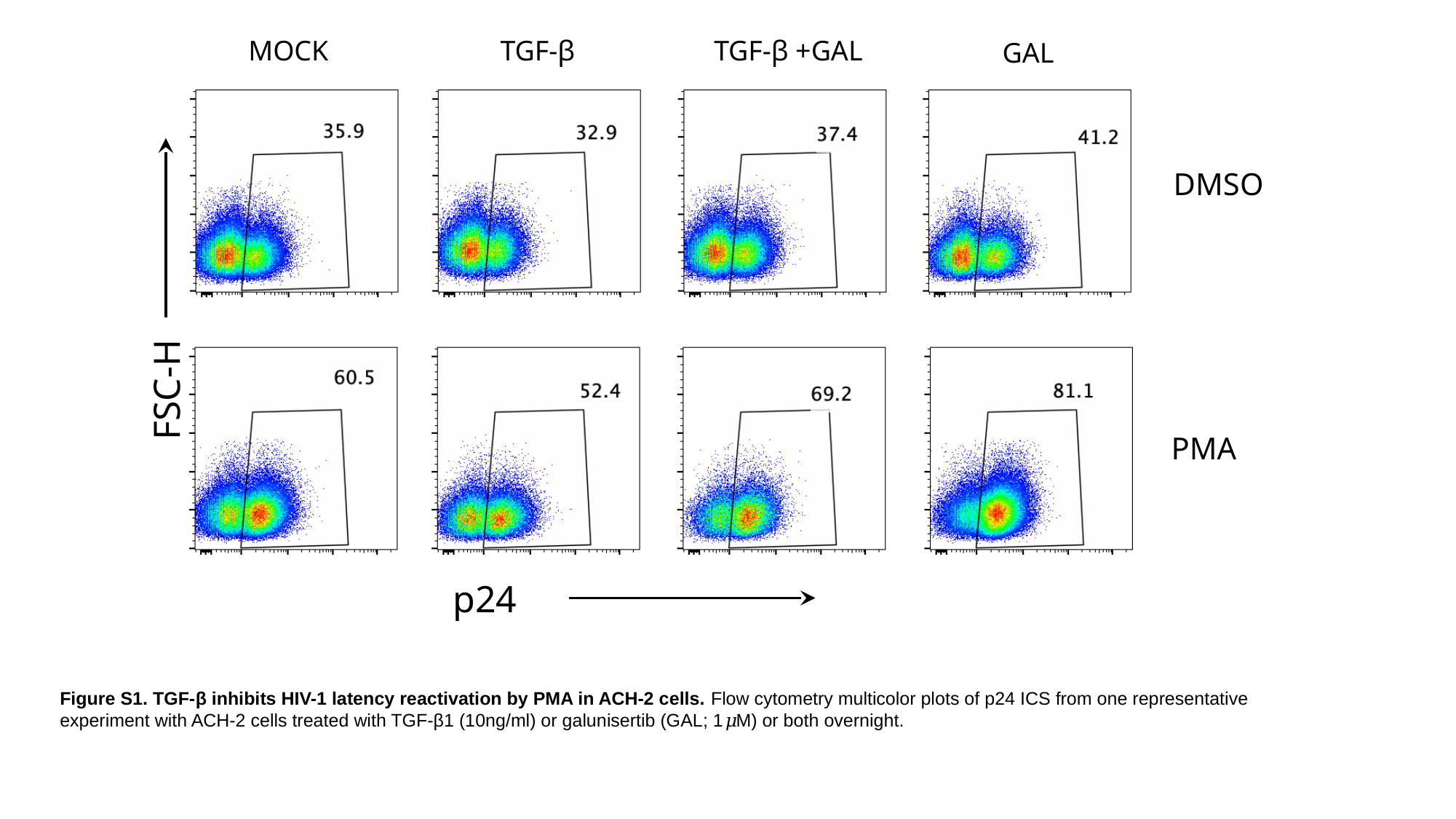

MOCK
TGF-β
TGF-β +GAL
GAL
DMSO
FSC-H
PMA
p24
Figure S1. TGF-β inhibits HIV-1 latency reactivation by PMA in ACH-2 cells. Flow cytometry multicolor plots of p24 ICS from one representative experiment with ACH-2 cells treated with TGF-β1 (10ng/ml) or galunisertib (GAL; 1𝜇M) or both overnight.

### Slide 2
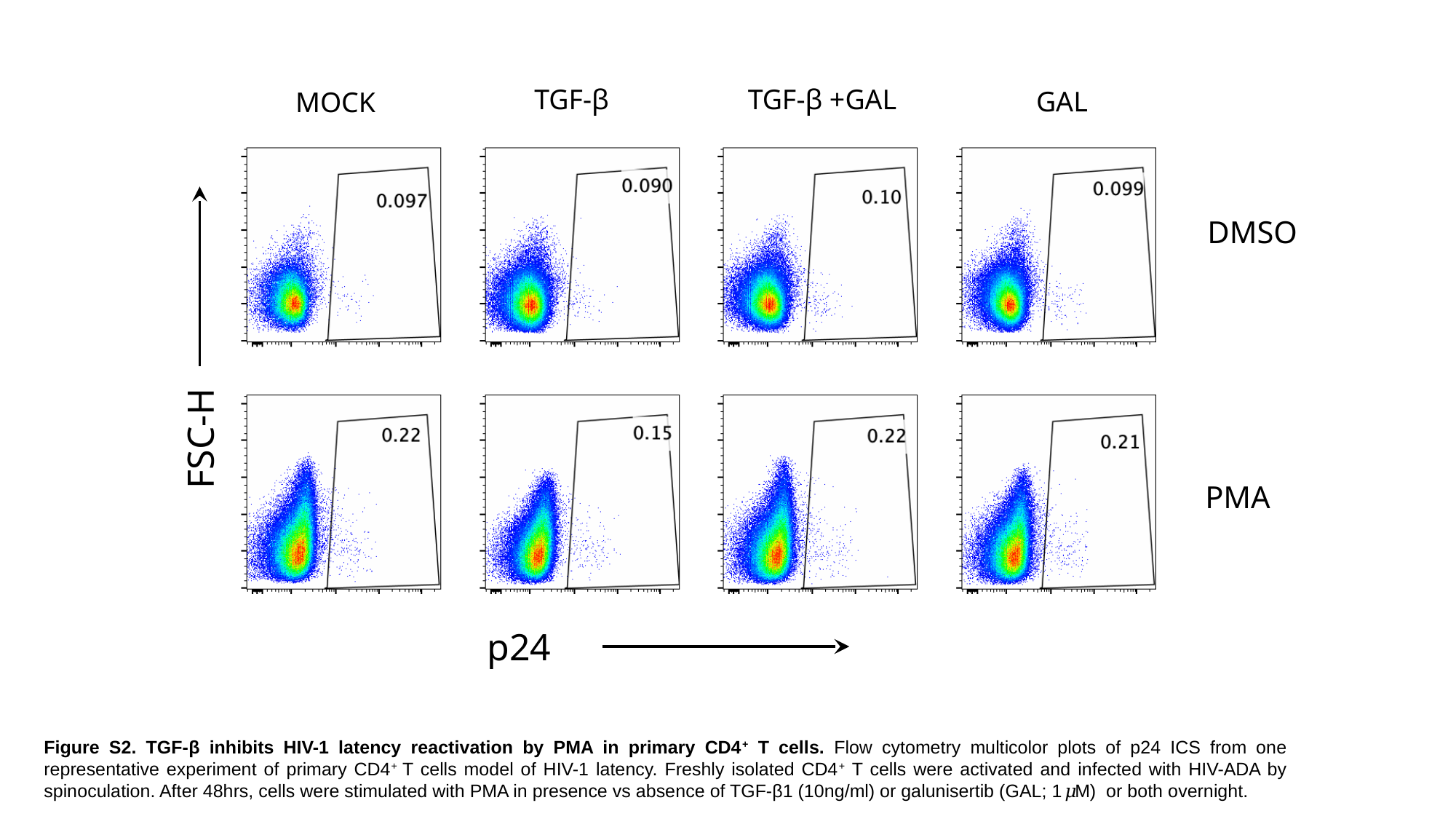

TGF-β
TGF-β +GAL
GAL
MOCK
DMSO
FSC-H
PMA
p24
Figure S2. TGF-β inhibits HIV-1 latency reactivation by PMA in primary CD4^+^ T cells. Flow cytometry multicolor plots of p24 ICS from one representative experiment of primary CD4^+^ T cells model of HIV-1 latency. Freshly isolated CD4^+^ T cells were activated and infected with HIV-ADA by spinoculation. After 48hrs, cells were stimulated with PMA in presence vs absence of TGF-β1 (10ng/ml) or galunisertib (GAL; 1𝜇M) or both overnight.

### Slide 3
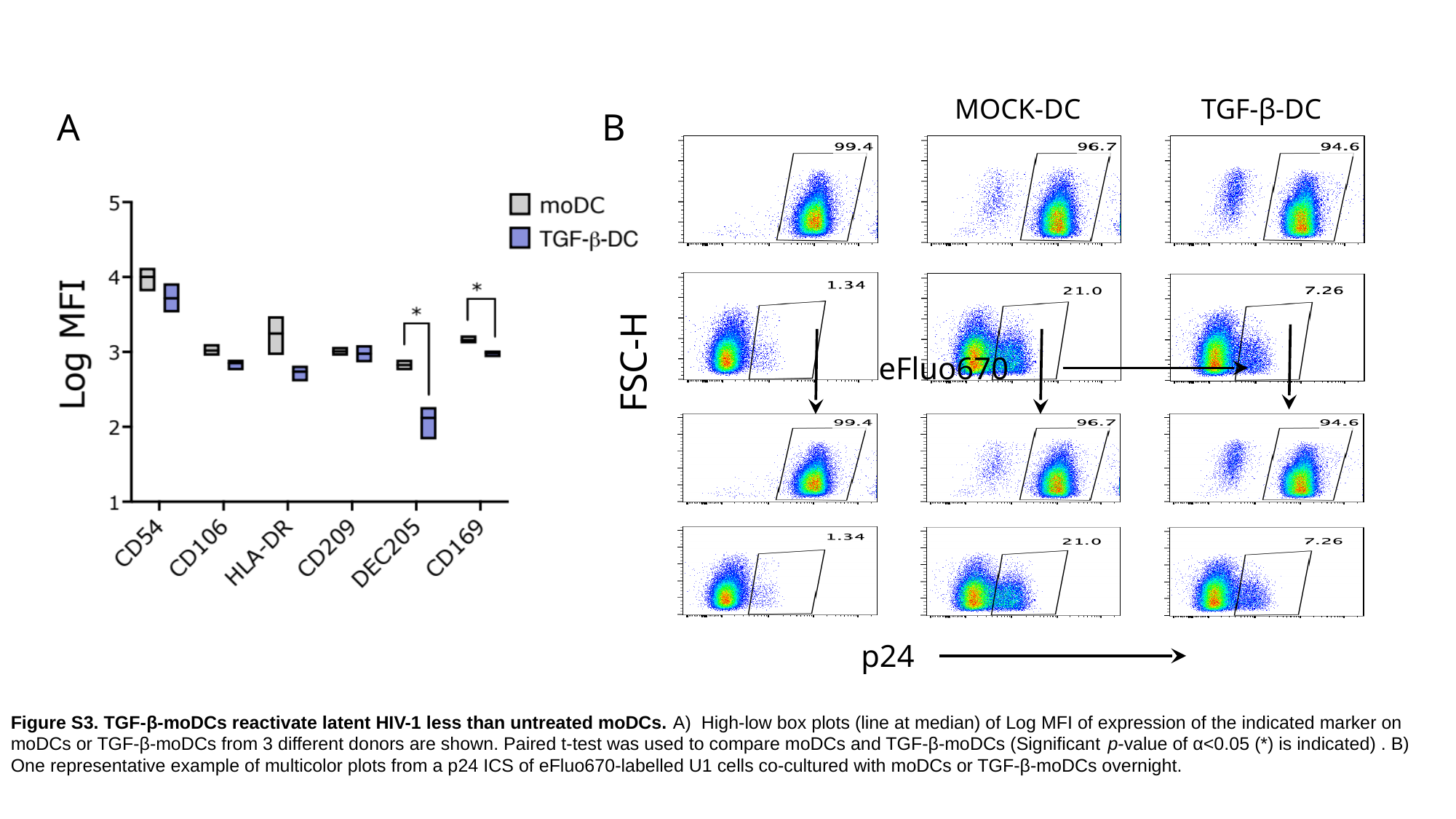

MOCK-DC
TGF-β-DC
A
B
FSC-H
eFluo670
p24
Figure S3. TGF-β-moDCs reactivate latent HIV-1 less than untreated moDCs. A) High-low box plots (line at median) of Log MFI of expression of the indicated marker on moDCs or TGF-β-moDCs from 3 different donors are shown. Paired t-test was used to compare moDCs and TGF-β-moDCs (Significant p-value of α<0.05 (*) is indicated) . B) One representative example of multicolor plots from a p24 ICS of eFluo670-labelled U1 cells co-cultured with moDCs or TGF-β-moDCs overnight.

### Slide 4
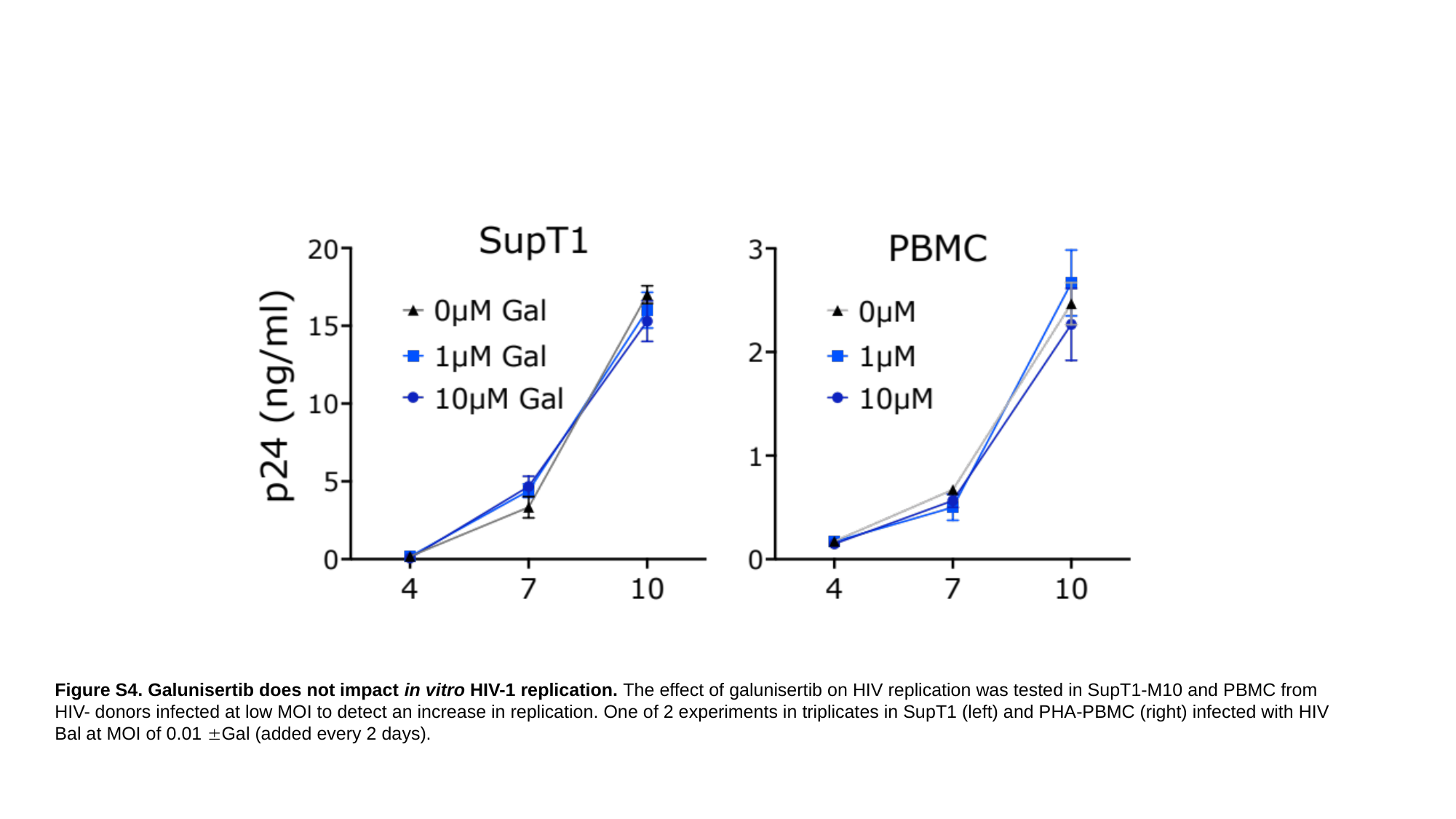

Figure S4. Galunisertib does not impact in vitro HIV-1 replication. The effect of galunisertib on HIV replication was tested in SupT1-M10 and PBMC from HIV- donors infected at low MOI to detect an increase in replication. One of 2 experiments in triplicates in SupT1 (left) and PHA-PBMC (right) infected with HIV Bal at MOI of 0.01 ±Gal (added every 2 days).

### Slide 5
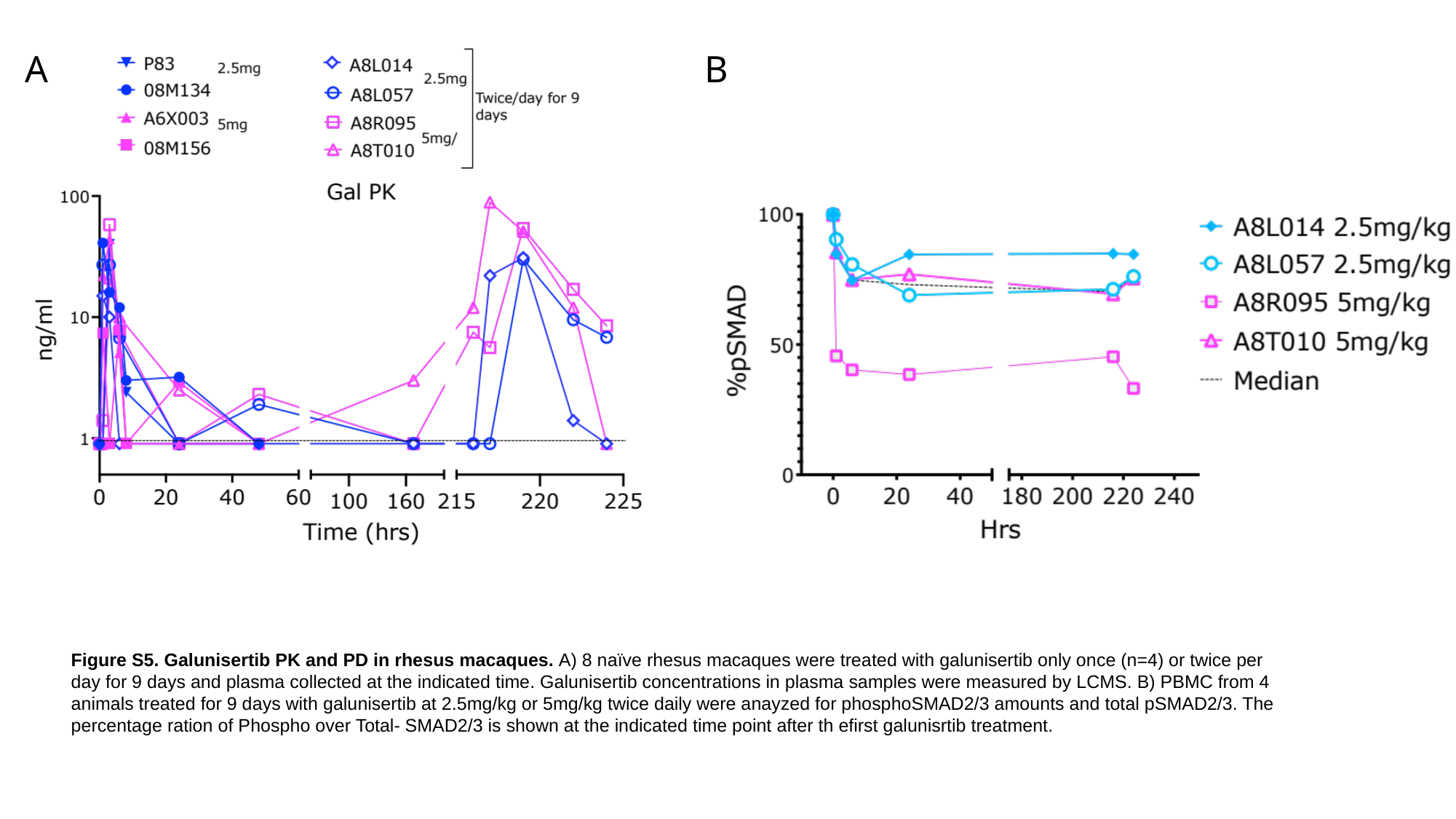

A
B
Figure S5. Galunisertib PK and PD in rhesus macaques. A) 8 naïve rhesus macaques were treated with galunisertib only once (n=4) or twice per day for 9 days and plasma collected at the indicated time. Galunisertib concentrations in plasma samples were measured by LCMS. B) PBMC from 4 animals treated for 9 days with galunisertib at 2.5mg/kg or 5mg/kg twice daily were anayzed for phosphoSMAD2/3 amounts and total pSMAD2/3. The percentage ration of Phospho over Total- SMAD2/3 is shown at the indicated time point after th efirst galunisrtib treatment.

### Slide 6
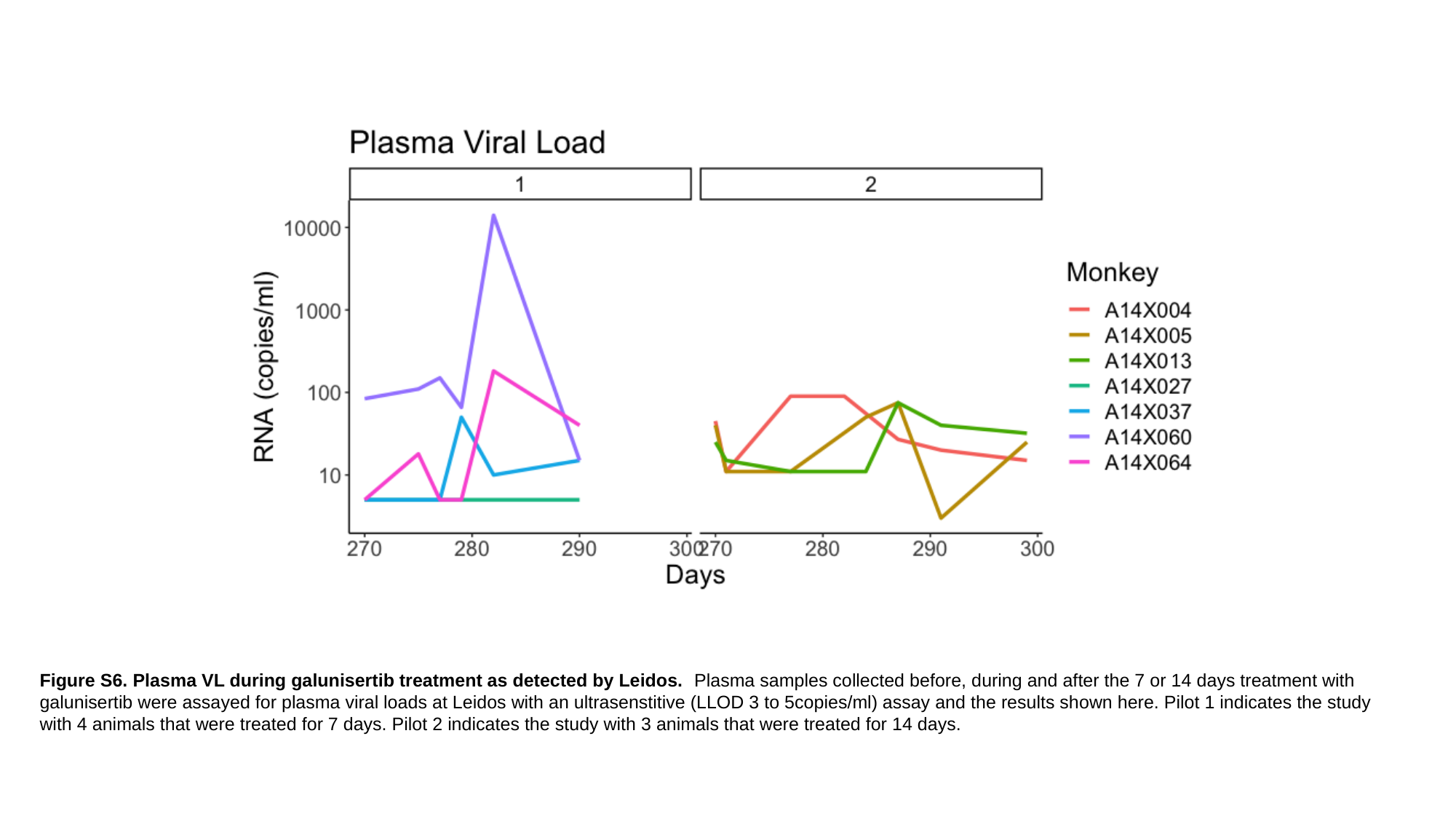

Figure S6. Plasma VL during galunisertib treatment as detected by Leidos. Plasma samples collected before, during and after the 7 or 14 days treatment with galunisertib were assayed for plasma viral loads at Leidos with an ultrasenstitive (LLOD 3 to 5copies/ml) assay and the results shown here. Pilot 1 indicates the study with 4 animals that were treated for 7 days. Pilot 2 indicates the study with 3 animals that were treated for 14 days.

### Slide 7
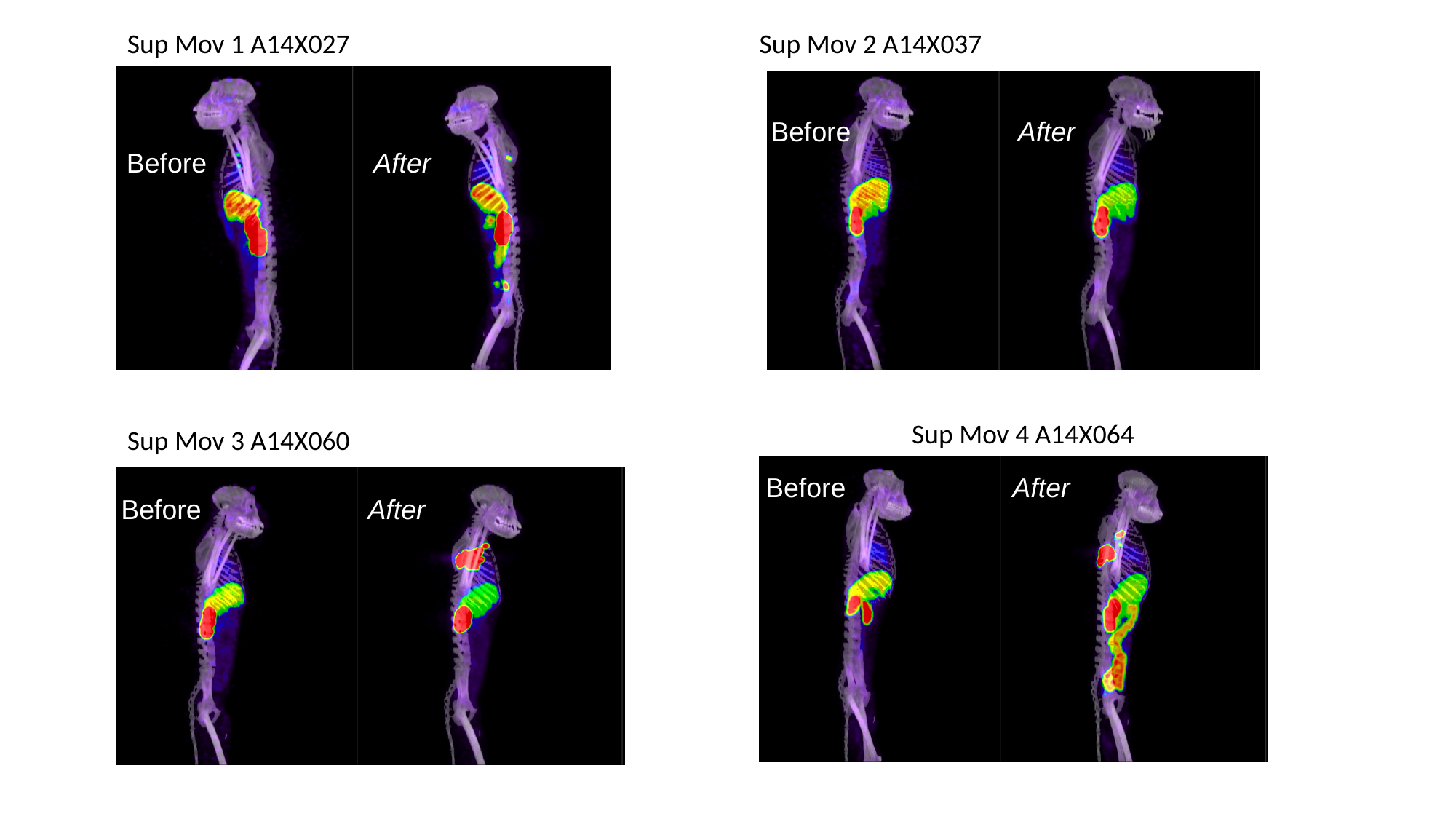

Sup Mov 2 A14X037
Sup Mov 1 A14X027
Before
After
Before
After
Sup Mov 4 A14X064
Sup Mov 3 A14X060
Before
After
Before
After

### Slide 8
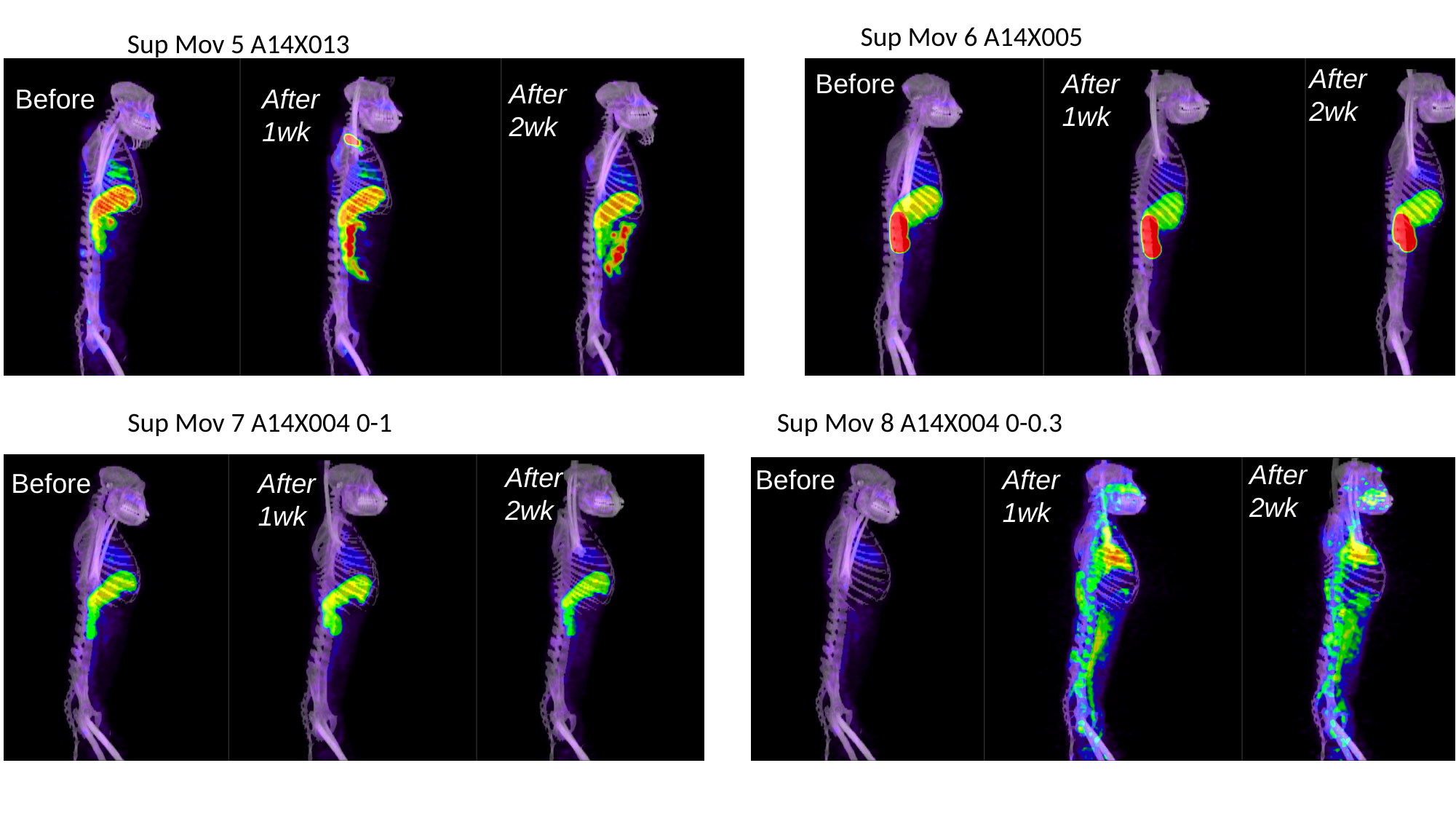

Sup Mov 6 A14X005
Sup Mov 5 A14X013
After
2wk
Before
After
1wk
After
2wk
Before
After
1wk
Sup Mov 7 A14X004 0-1
Sup Mov 8 A14X004 0-0.3
After
2wk
After
2wk
Before
After
1wk
Before
After
1wk

### Slide 9
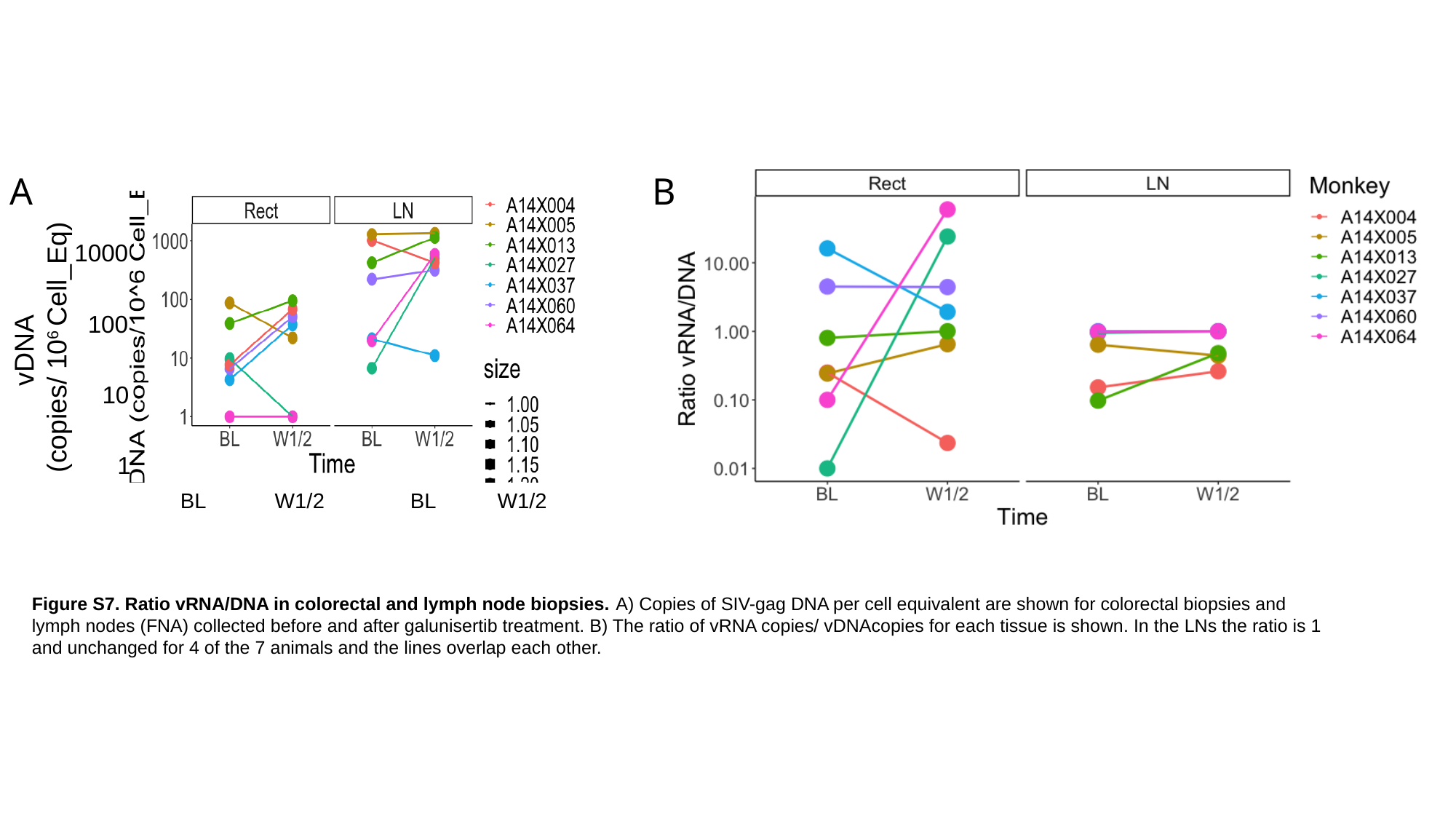

A
B
1000
100
10
1
BL
W1/2
BL
W1/2
vDNA
(copies/ 10^6^ Cell_Eq)
Figure S7. Ratio vRNA/DNA in colorectal and lymph node biopsies. A) Copies of SIV-gag DNA per cell equivalent are shown for colorectal biopsies and lymph nodes (FNA) collected before and after galunisertib treatment. B) The ratio of vRNA copies/ vDNAcopies for each tissue is shown. In the LNs the ratio is 1 and unchanged for 4 of the 7 animals and the lines overlap each other.

### Slide 10
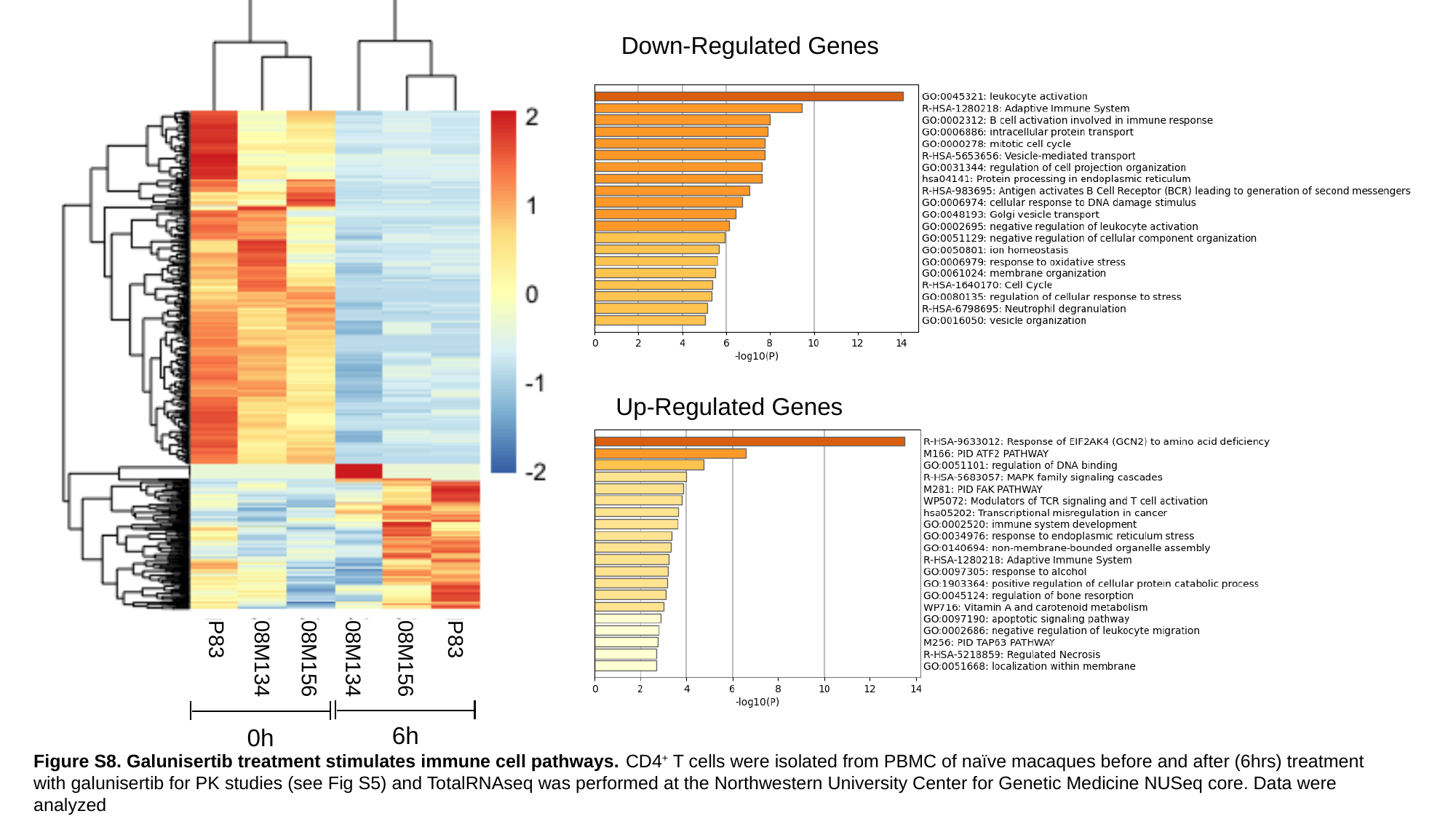

Down-Regulated Genes
Up-Regulated Genes
P83
P83
08M156
08M156
08M134
08M134
6h
0h
Figure S8. Galunisertib treatment stimulates immune cell pathways. CD4^+^ T cells were isolated from PBMC of naïve macaques before and after (6hrs) treatment with galunisertib for PK studies (see Fig S5) and TotalRNAseq was performed at the Northwestern University Center for Genetic Medicine NUSeq core. Data were analyzed

### Slide 11
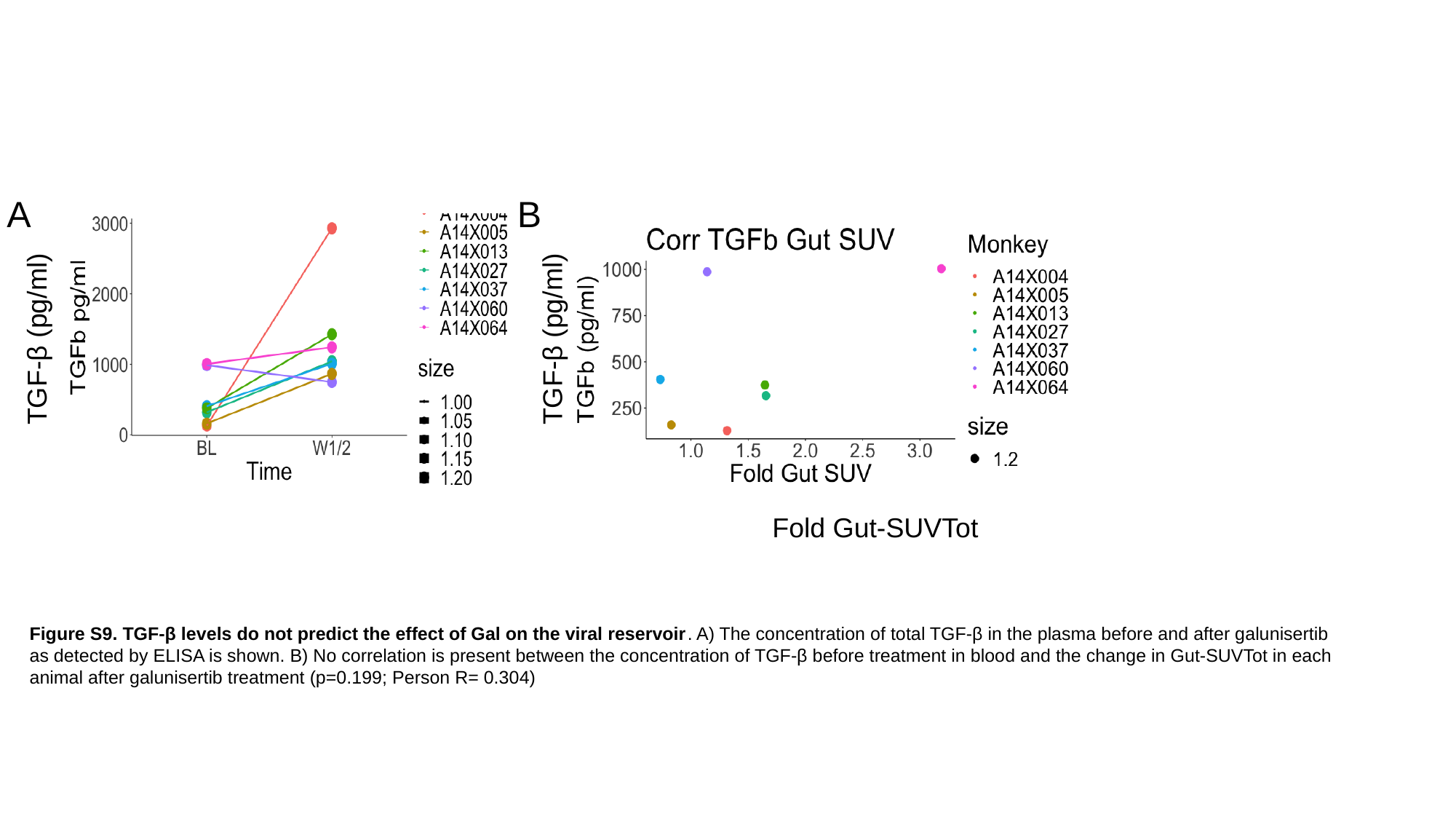

A
B
TGF-β (pg/ml)
TGF-β (pg/ml)
Fold Gut-SUVTot
Figure S9. TGF-β levels do not predict the effect of Gal on the viral reservoir. A) The concentration of total TGF-β in the plasma before and after galunisertib as detected by ELISA is shown. B) No correlation is present between the concentration of TGF-β before treatment in blood and the change in Gut-SUVTot in each animal after galunisertib treatment (p=0.199; Person R= 0.304)
